# Environmental DNA reveals fine scale spatial and temporal variation of prey species for marine mammals in a Scottish marine protected area

**DOI:** 10.1101/2023.12.21.572838

**Authors:** Elizabeth Boyse, Kevin P. Robinson, Maria Beger, Ian M. Carr, Morag Taylor, Elena Valsecchi, Simon J. Goodman

## Abstract

Marine mammal foraging grounds are popular focal points for marine protected area (MPA) implementation, but may be temporally dynamic, requiring continuous monitoring to infer prey availability and abundance. Marine mammal distributions are assumed to be driven by their prey in foraging areas, but limited understanding of prey distributions often prevents us from exploring how shifting prey availability impacts both seasonal and long-term marine mammal distributions. Environmental DNA (eDNA) metabarcoding could enhance understanding of marine mammal habitat use in relation to their prey through simultaneous monitoring of both. However, eDNA applications focused on marine mammals or predator-prey dynamics have been limited to date. In this study, we assess spatiotemporal changes in the availability and abundance of minke whale (*Balaenoptera acutorostrata*) prey species in a newly established MPA, employing eDNA metabarcoding. We recovered 105 molecular operational taxonomic units (OTUs) from marine vertebrates using two primer sets targeting 12S and 16S genes, along with 112 OTUs from a broader eukaryotic primer set targeting 18S rRNA. Overall, key forage fish prey species, sandeels and clupeids, were the most abundant teleost fishes detected, although their availability varied temporally and with distance from shore. We also found clear spatial partitioning between coastal bottlenose dolphins and the more pelagic minke whales and harbour porpoises, paralleling availability of their main prey species. Other species of conservation interest were also detected including the critically endangered European eel (*Anguilla anguilla*), blue fin tuna (*Thunnus thynnus*), and the invasive pink salmon (*Oncorhynchus gorbuscha*). This study demonstrates the application of eDNA to detect spatiotemporal trends in the occurrence and abundance of cetacean predators and their prey, furthering our understanding of fine-scale habitat use within MPAs. Future, long-term monitoring of predator-prey dynamics with eDNA could improve our ability to predict climate-induced shifts in foraging grounds and enhance rapid responses with appropriate management actions.

## Introduction

Temperate nearshore marine habitats experience high levels of anthropogenic impacts and are predicted to experience significant ecological change as a result of climate heating (O’Hara et al., 2021; Williams et al., 2022). There is an increasing need to expand spatiotemporal capacity for biodiversity monitoring to track the status of ecosystems and individual species, and to help evaluate vulnerability and exposure to anthropogenic activities (McQuatters-Gollop et al., 2022). Here we test the capacity for environmental DNA (eDNA) metabarcoding to resolve fine scale seasonal and spatial variation in ecological community structure for vertebrate and invertebrate taxa, and patterns of habitat use by keystone cetacean species in a newly designated marine protected area in the Moray Firth, Scotland.

Over a quarter of all European marine mammal species are threatened as a result of overfishing, shipping traffic, pollution, changes in prey dynamics and habitat degradation (Avila et al., 2018; Braulik et al., 2023), jeopardising the important ecosystem functions marine mammals provide (Estes et al., 2016). Marine protected areas (MPAs) are the main tool used to protect marine mammals from human impacts. However, European MPAs are currently too small and disjointed to provide adequate protection for such highly mobile and far ranging mammals (Bearzi & Reeves, 2021), with complex, dynamic, seasonal partitioning of foraging and breeding, sensitive to long term environmental change (Notarbartolo Di Sciara et al., 2016).

Drivers of cetacean distributions are assumed to have a hierarchical structure, with distributions at fine spatiotemporal resolutions (10s of kilometres) best described by prey availability, while at broader scales (100s of kilometres), oceanographic features become more important (Mannocci et al., 2017). However, prey data are rarely available at complementary spatiotemporal scales to parameterise predictive distribution models for marine mammals, so environmental proxies are used instead (Mannocci et al., 2020; Pendleton et al., 2020). Improved understanding of the relationship between cetaceans and their prey will be vital to accurately predict future distribution shifts as both respond to climatic change, especially to mitigate against species moving into unprotected areas or areas with higher exposure to threats (Agardy et al., 2019; Silber et al., 2017).

Recent advances in environmental DNA metabarcoding offer the opportunity to simultaneously monitor cetaceans and their prey, enabling long-term tracking of distributions, and to enhance our understanding of their dynamics (Székely et al., 2021). eDNA can expand the spatiotemporal scope of marine mammal monitoring where visual or acoustic monitoring are infeasible, *i.e.*, at night or in adverse weather conditions and for cetaceans that vocalise infrequently or have unknown vocalisations (Baumgartner et al., 2019; Valsecchi et al., 2021). To date, single species eDNA approaches have improved our understanding of several rare or threatened species, *i.e.*, dwarf sperm whales (*Kogia sima*) and Mediterranean monk seals (*Monachus monachus*) (Juhel et al., 2021; Valsecchi et al., 2023), and species that are challenging to detect with conventional survey techniques, *i.e.*, beaked whales (Boldrocchi et al., 2023; Hooker et al., 2019). They have also provided insights into population genetics, with important management consequences (Parsons et al., 2018). Dietary metabarcoding studies have uncovered previously unknown marine mammal diets (Sonsthagen et al., 2020; Zhang et al., 2023), whilst eDNA metabarcoding has revealed spatiotemporal availability of prey species and detected co-occurrences between cetaceans and their prey (Djurhuus et al., 2020; Visser et al., 2021). However, few studies have harnessed eDNA to elucidate marine mammal trophic interactions to date (Székely et al., 2021).

The Southern Trench MPA in the Moray Firth, north-east Scotland, has recently been designated to protect important seasonal foraging grounds for minke whales (*Balaenoptera acutorostrata*) (NatureScot, 2020). The diet of this baleen whale within the Moray Firth consists predominantly of sandeels (*Ammodytes* sp.), sprat (*Sprattus sprattus*) and herring (*Clupea harengus*), none of which are commercially fished in the area, resulting in limited knowledge of their spatiotemporal availability and abundance (Pierce et al., 2004). The area also overlaps with bottlenose dolphin (*Tursiops truncatus*) and harbour porpoise (*Phocoena phocoena*) distributions, which require protection under Annex II of the European Habitats Directive. Harbour porpoises also rely predominantly on sandeels as well as whiting (*Merlangius merlangus*), whilst bottlenose dolphins target Gadidae species including cod (*Gadus morhua*), saithe (*Pollachius virens*) and whiting, and salmonids (*Salmo* sp.) (Santos et al., 2004; Santos et al., 2001). In this study, we characterise the vertebrate and broader eukaryotic community composition within the MPA, and assess temporal changes across the minke whale foraging season (June to October), to evaluate the strength of seasonality in prey availability. We further examine how the community changes with distance from the shore, to explore how prey distributions influence the distribution of sympatric cetacean species with different dietary preferences. We expect that seasonal changes in minke whale habitat use correlate with spatiotemporal changes in the availability of different prey species, potentially indicating prey switching. The work provides a monitoring baseline for this newly established MPA and a foundation to develop longer-term eDNA monitoring protocols that improve ecological knowledge and contribute to the adaptive management process. It also highlights the potential for similar monitoring approaches to be applied to MPAs supporting cetaceans and other large marine predators worldwide.

## Methods

### Study Area

At 5,230 km^2^, the Moray Firth (Figure 1) is the largest estuarine embayment in northeast Scotland, and an extension of the North Sea basin beyond (Harding-Hill, 1993). It is an internationally recognised area of outstanding biological importance, with the ‘inner’ region designated as a Special Area of Conservation (SAC) under the European Habitats Directive (92/43/EEC) (Cheney et al., 2013), as well as the newly designated Southern Trench MPA in the south-eastern ‘outer’ firth (NatureScot, 2020). The Southern Trench is an enclosed basin reaching depths in excess of 250 m, constituting the deepest portion of the firth (Holmes et al., 2004). Two oceanographic features govern water movement. Firstly, cold water is transported into the firth from the northern North Sea via the Dooley current, whilst a warm water plume ebbs out from the inner firth and rivers discharging into the firth, which is associated with greater primary productivity levels (Tetley et al., 2008).

**Figure 1.**
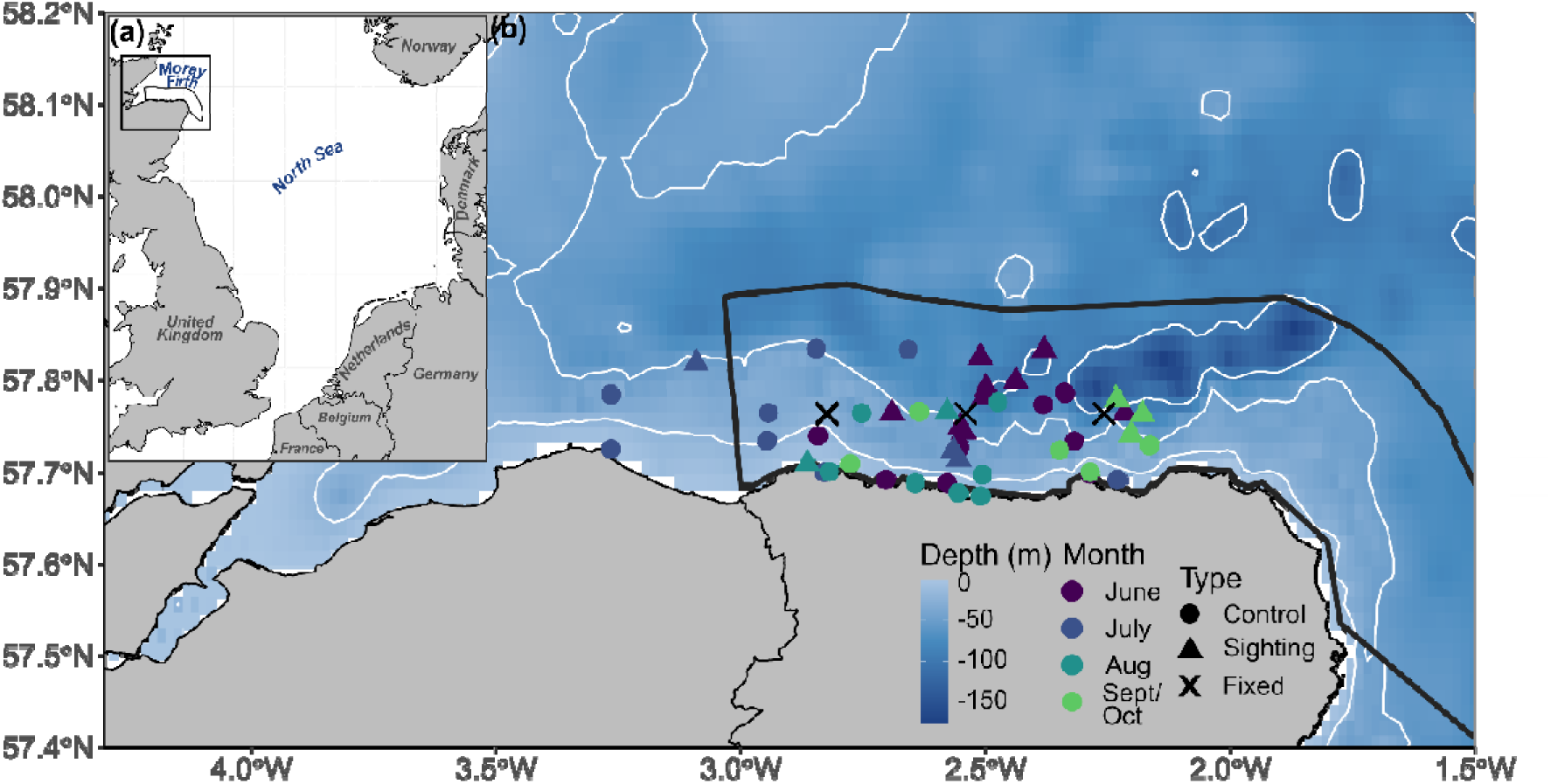
Maps showing (a) the location of the Moray Firth within the North Sea; and (b) the position of the Southern Trench MPA (black outline polygon). The positions of the fixed eDNA sampling points are illustrated with crosses, with control and sightings samples indicated by circles and triangles, coloured by sampling month.

### Sample Collection

Seawater samples were collected from June to October 2021, corresponding to the months when minke whales are most abundant within the firth (Robinson et al., 2009). Samples were collected on 18 different sampling days across four monthly sampling trips to capture spatiotemporal trends in the presence and relative abundance of the main whale prey species and overall community trends (Appendix Table A1). All sampling was carried out from an 8-metre rigid hulled inflatable boat (RHIB) using a weighted bucket deployed to a depth of four metres. Eleven litres of the resulting sea water sample was subsequently transferred to sealed sterile plastic-aluminium ‘Bags in the Box’ containers for storage and transport to the laboratory (Valsecchi et al., 2021). Reusable field equipment was cleaned with 50% bleach, left to soak for 30 minutes and then washed thoroughly with tap water between sampling trips.

To quantify temporal differences in community composition, samples were collected from three fixed positions (Figure 1), located five nautical miles offshore, yielding 12 fixed samples over the 4 months. The bathymetry of the westerly fixed sampling point was the shallowest, at 33 m, whilst the easterly fixed point was on the edge of the Southern Trench at 118 m. The middle-fixed point was at 104 m depth, and a known hotspot for foraging minke whales. In addition, samples were collected whenever minke whales were sighted on the eDNA sampling days. This was to assess whether whales needed to be in close proximity to detect their DNA, and the extent of co-detection with prey species (total 18 sightings samples). Finally, random samples were collected across the entire study area (total 30 control samples) to facilitate evaluation of the environmental drivers of community composition (Figure 1). This design resulted in a total of 57 11-litre seawater samples, plus three field-controls (blanks) which were collected in July, August and September with the same sampling equipment but replacing seawater with tap water to detect potential sources of contamination.

### Sample Filtration

We filtered samples between one and six days after collection, with an average delay of 1.8 ± 1.1 s.d. days. 49% of samples were filtered the day after collection, and 89% of samples were filtered within three days of collection. Each 11-litre seawater sample was split into three replicates for filtering: two 4-litre replicates and one 3-litre replicate. For ten of the samples, between 10 and 10.8 litres was filtered as a result of filters being saturated before 11 litres had been reached. Samples were filtered using either the BioSart^®^ 100 filtration system (Sartorius) or Nalgene^TM^ reusable analytical funnels (Thermo Scientific) with either the Fisherbrand^TM^ FB70155 Pump or Welch^TM^ WOB-L Piston Dry Vacuum Pump. Samples were filtered through cellulose nitrate filters of 0.45 μm pore size. Immediately after filtration, the filter papers were wrapped and stored in aluminium foil at −20°C.

### DNA extraction, amplification and sequencing

DNA extractions were carried out in a dedicated molecular laboratory, and bench surfaces were cleaned with 50% bleach followed by deionised water. Extractions were carried out using a Qiagen DNeasy PowerSoil Pro Kit following the manufacturer’s protocol.

Pre- and post-PCR procedures were carried out in separate rooms. All PCRs were prepared in a class II microbiology safety cabinet that was cleaned with 50% bleach and illuminated with ultraviolet light for 20 minutes between sample preparations. We amplified marine vertebrate DNA with two primer sets: MarVer1, which amplifies an approximately 202 bp sequence from the mitochondrial 12S rRNA gene; and MarVer3, which amplifies an approximately 245 bp sequence from the mitochondrial 16S rRNA gene (Valsecchi et al., 2020). These primers can resolve most taxa to species-level across all marine vertebrates, inclusive of marine mammals, elasmobranchs and teleost fishes. Our primers were designed with six to eight random nucleotides, an eight-base pair Illumina barcode tag and the amplification primer sequence from 5’ to 3’ (Bohmann et al., 2022). PCR reactions were 20 μL volume containing 0.025 u/μL GoTaq® Hot Start Polymerase (Promega), 5X Green GoTaq® Flexi Buffer (Promega), 1mM or 2mM MgCl_2_ (Promega) for MarVer1 or MarVer3 respectively, 0.2 mM dNTPs (Promega), 0.2 μM each of the reverse and forward primer, and UltraPure^TM^ distilled water (Invitrogen). Annealing temperatures for MarVer1 were 54/55/56°C for 10/10/18 cycles, and 58/57/56/55°C for 8/10/10/10 cycles for MarVer3. Both had an initial denaturation time of four minutes at 95°C, and a final elongation of five minutes at 72°C, then per cycle, 30 seconds at 95°C, 10 s at annealing temperature and 20 s at 72°C for MarVer1; and for MarVer3, 30 s at 95°C, 2 s for the first 8 cycles and 10 s for remaining cycles at annealing temperature, and 20 s at 72°C. Three PCR replicates were amplified per 11-litre water sample for each primer set, and then pooled for individual samples per primer set. Amplicons were cleaned up and primer dimers removed with AMPure beads (Beckman Coulter). We then checked that the fragment size distributions were as expected with an Agilent TapeStation, followed by quantification with a Qubit fluorometer. Amplicons for each primer set were pooled in equimolar ratios to create two Illumina NEBNext Ultra DNA libraries, one for MarVer1 and MarVer3 respectively. The libraries were sequenced separately on an Illumina MiSeq Sequencer with 150 bp paired-end lanes at the University of Leeds Genomics Facility, St James’s Hospital.

We also amplified an approximately 260 bp amplicon for the V9 region of 18S rRNA using a general eukaryotic primer set, 1391F and EukBr, to detect zooplankton and other invertebrates (Amaral-Zettler et al., 2009). Primers included sequences homologous to Illumina sequencing adapters appended to the 5’ end. PCR reactions were 25 μL, consisting of 0.025 u/μL GoTaq® Hot Start Polymerase (Promega), 5X Green GoTaq® Flexi Buffer (Promega), 2 mM MgCl_2_ (Promega), 0.2 mM dNTPs (Promega), 0.2 μM each of the reverse and forward primer, 1.6 μM mammal blocking primer, and UltraPure^TM^ distilled water (Invitrogen). Thermocycling conditions included an initial denaturation at 94°C for three minutes, 35 cycles at 94°C for 45 s, 65°C for 15 s, 57°C for 30 s, 72°C at 90 s, and a final elongation at 72°C for ten minutes (Sawaya et al., 2019). The three PCR replicates per 11-litre water sample were then pooled and cleaned up as above. The final sequencing libraries were generated using a second PCR in which Nextera XT indexed adaptor sequences were used as primers, such that each sample was uniquely indexed. The PCR reaction consisted of 5 μL of the pooled amplicons, 25 μL of the NEBNext Q5 Hot Start HiFi PCR Master Mix, 10 μL of water and 5 μL each of the appropriate Nextera XT Index Primer 1 and Primer 2. Thermocycling conditions included an initial denaturation at 95°C for three minutes, followed by 8 cycles of 95°C for 30 s, 55°C for 30 s, 72°C for 30 s and a final hold at 72°C for five minutes. The products were then cleaned again with AMPure beads to remove adapter dimers and free adaptor oligos and checked for the presence of the correctly formed libraries by running on a D1000 tape of a TapeStation followed by quantification with Qubit fluorometry. The libraries were then combined to create an equimolar pool that was sequenced on a MiSeq 250 bp paired end lane with V2.0 chemistry and 15% PhiX control library to aid base calling.

### Bioinformatics

A description of the full bioinformatics workflow is available in Valsecchi et al. (2020) and at http://www.dna-leeds.co.uk/eDNA/. In brief, initial quality checking was performed to remove read pairs with spurious primer combinations and trimmed to remove low quality sequences, before read pairs were combined to form a single sequence. Only one occurrence of each unique sequence per sample was retained to reduce the likelihood of PCR duplicates or chimaeras. A counts matrix was created by aggregating the number of instances for each unique amplicon sequence per sample. Amplicon sequences were blasted against the Genbank Nucleotide Database to identify the taxonomic origin of sequences, and the top ten hits were linked to the sequence if they were more than 70% homologous. Full taxonomic hierarchy for species names was obtained for the GenBank hit sequences from a Microsoft SQL server instance of the ITIS taxonomy database. When taxonomic information was found, the name and taxonomy of the best hit was retained. Finally, sequences were clustered into molecular operational taxonomic units (OTUs) using a 98% threshold of homology to the GenBank hit sequence at species level for vertebrate primer sets, and a 95% threshold of homology at family level for our 18S primer set.

### Contamination control

For the purpose of this study, non-marine OTUs and off target OTUs, *i.e.,* invertebrates detected with our vertebrate primer sets, were removed. Non-marine OTUs were primarily comprised of *Homo sapiens* and agricultural species such as *Bos taurus* and *Sus scrofa, Canis lupus,* local terrestrial wildlife including *Capreolus capreolus* (roe dear), *Erinaceus europaeus* (hedgehog) and *Myodes glareolus* (voles). Given the coastal nature of our study area, these detections potentially originate from river inflow and other terrestrial water runoff (sewage, storm drains etc). For our vertebrate primers, we identified amplicon sequences which had been assigned to marine species not previously recorded in the North Sea, according to FishBase records (www.fishbase.org). We established whether native congenerics or family members were known to be present in the North Sea, and if so, compared amplicon sequences to assess whether there was enough differentiation in our amplicon regions to distinguish between the non-indigenous and native congenerics or family members (Valsecchi et al., 2021). Non-native species reads were either confirmed as a potential invasive species, reassigned to a native congeneric or family group, or excluded if they had fewer than 10 reads. A full list of species that were removed or merged is provided in the appendix (Appendices A3 and A4 respectively).

The most likely source of contamination in this study was from crossover contamination between samples (Calderón□Sanou et al., 2020). To reduce this background contamination, we calculated two times the standard deviation for the proportion of each OTU in the field blanks, and then subtracted this from the specific OTU proportion in each sample, following a similar approach to Kelly et al. (2018). To facilitate abundance comparisons, we standardised read counts using an OTU-specific index under the assumption that amplification efficiency is consistent for each OTU, regardless of community composition. This scaling allows for the comparison of within OTU abundance across samples (Kelly et al., 2019). The OTU-specific index was made by converting read counts into proportions then dividing the maximum proportion for every OTU from that OTU’s proportion per sample, resulting in an index between 0-1 for each OTU (Kelly et al., 2019). For our vertebrate primer sets, we also created an ensemble OTU index by averaging across the indices per sample for each of the primer sets (Kelly et al., 2019).

### Community analysis

Initial descriptions of community composition and visualisations of the data were conducted using the ‘Phyloseq’ R package version 1.38.0 (McMurdie & Holmes, 2013). We analysed temporal trends in the community through partitioning the data by sampling month, and spatial trends by dividing the samples into categories based on their distance from shore; near (<1.2 km), middle-near (between 2.5 and 7 km), middle-far (between 7 and 10 km), and far (> 10 km up to 16.1 km) (Drummond et al., 2021; Fraija□Fernández et al., 2020). These categories were defined based on previous observations that juvenile minke whales were most frequently sighted inshore (<2.5 km), whilst adults more frequently occurred further offshore (>10 km) (Robinson et al., 2023).

Community statistics were calculated using the ‘Vegan’ R package version 2.5-7, and using abundance data with the OTU-specific index applied (Dixon, 2003). Alpha diversity of samples was estimated with the Shannon-Weiner index and compared across the sampling months and with distance from shore using Kruskal-Wallis tests, followed by pairwise Wilcoxon rank sum tests. We transformed our abundance matrix into a Bray-Curtis dissimilarity matrix to compare beta-diversity between sampling months and distance from shore categories. We assessed differences between communities with non-metric multidimensional scaling (NMDS) and tested for significant differences between communities using permutational multivariate analysis of variance (PERMANOVA). Homogeneity of group dispersion is an assumption of PERMANOVA, so this was first assessed using the ‘betadisper’ function. We also performed pairwise multilevel comparisons to further evaluate where differences between communities existed using the ‘PairwiseAdonis’ R package version 0.4 (Martinez Arbizu, 2017). We identified indicator species for the different months and distance from shore categories using the ‘multipatt’ function, with 999 permutations, from the ‘Indicspecies’ R package version 1.7.12 (Cáceres & Legendre, 2009). This included two components: (A) which quantifies the specificity of the species as an indicator for the group where A=1 means that species only appears in that group, and (B) which quantifies the sensitivity of the species as an indicator for that group where B = 1 means that all sites within that group include the species (Cáceres & Legendre, 2009).

We evaluated environmental covariates associated with changes in community composition using multiple regression in distance matrices (MRM), employing Spearman correlation ranked distances and 10,000 permutations with the ‘MRM’ function from the ‘Ecodist’ R package version 2.0.9 (Goslee & Urban, 2007). We used four environmental predictors, bathymetry (m), sea surface temperature (SST; °C), chlorophyll *a* concentration (mg/m^3^) and distance from shore (m). Bathymetry was obtained from the General Bathymetric Chart of the Oceans (GEBCO) at a 0.004×0.004° resolution (GEBCO Compilation Group, 2020), and SST (https://doi.org/10.48670/moi-00156) and chlorophyll *a* (https://doi.org/10.48670/moi-00289) were downloaded from the E.U. Copernicus Marine Service Information at 0.02°x0.02° and 0.01°x0.01° resolutions respectively (Høyer & Karagali, 2016). Values for each predictor were then extracted from individual sampling points. Distance from shore was calculated using the ‘dist2Line’ function from the ‘Geosphere’ R package version 1.5-14 (Hijmans, 2021). Prior to running MRM, we tested for collinearity among predictor variables with Spearman’s rho rank correlation coefficient from the ‘Hmisc’ R package version 4.7-1 (Harrell Jr, 2022). Bathymetry and distance from shore were highly correlated (rho = −0.81, p < 0.001) so only distance from shore was retained for MRM. We created distance matrices for each environmental predictor using Euclidean distance, as well as a distance matrix for the distance between sites, and the number of days between sample collection. Our response variable was a Bray-Curtis dissimilarity matrix of community composition, derived from either our vertebrate ensemble OTU index or eukaryotic OTU index. Initially, maximal models were created using all terms, and then reduced to a minimal model with only significant terms retained.

## Results

### Composition of vertebrate taxa detected

Following contamination removal procedures, our final datasets for vertebrate primers MarVer1 and MarVer3 comprised 2, 880, 775 and 4,013,997 sequences which were assigned to 56 and 80 operational taxonomic units (OTUs) respectively, clustered at species level where possible (Table 1). 31 OTUs were detected by both markers, whilst 25 were unique to MarVer1, and 49 to MarVer3 (Figure 2). Over 90% of the reads from both markers were assigned to teleost fishes, from 22 families for MarVer1 and 31 families for MarVer3 (20 shared families, 2 unique to MarVer1 and 11 unique to MarVer3) (Figure 3a). This list included species of potential conservation interest, such as rare taxa (bluefin tuna, *Thunnus thynnus*), critically endangered species (European eel, *Anguilla anguilla*) and invasive species (pink salmon, *Oncorhynchus gorbuscha*) (Appendix Table A1). Mammalia had the second highest proportion of reads, including all four marine mammals common to the study area, namely the minke whale (*Balaenoptera acutorostrata*), grey seal (*Halichoerus grypus*), harbour porpoise (*Phocoena phocoena*), and bottlenose dolphin (*Tursiops truncatus*), as well as some less commonly sighted vagrants including the fin whale (*Balaenoptera physalus* – MarVer1 only), white-beaked dolphin (*Lagenorhynchus albirostris*-MarVer3 only) and Sowerby’s beaked whale (*Mesoplodon bidens*).

**Figure 2.**
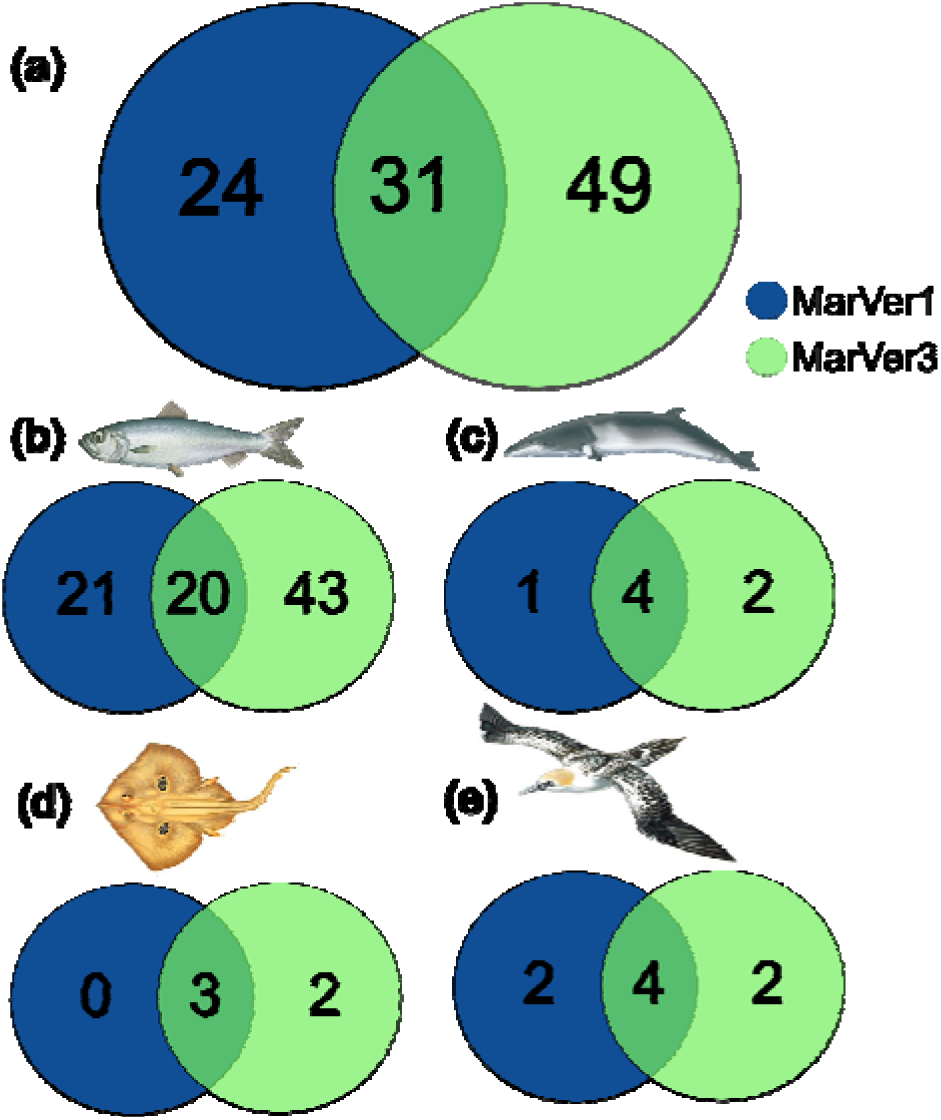
Venn diagrams showing the overlap between (a) all OTUs, (b) teleosts, (c) mammals, (d) chondrichthyes and (e) birds, detected by both vertebrate primer sets, MarVer1 (blue) and MarVer3 (green) respectively.

**Figure 3.**
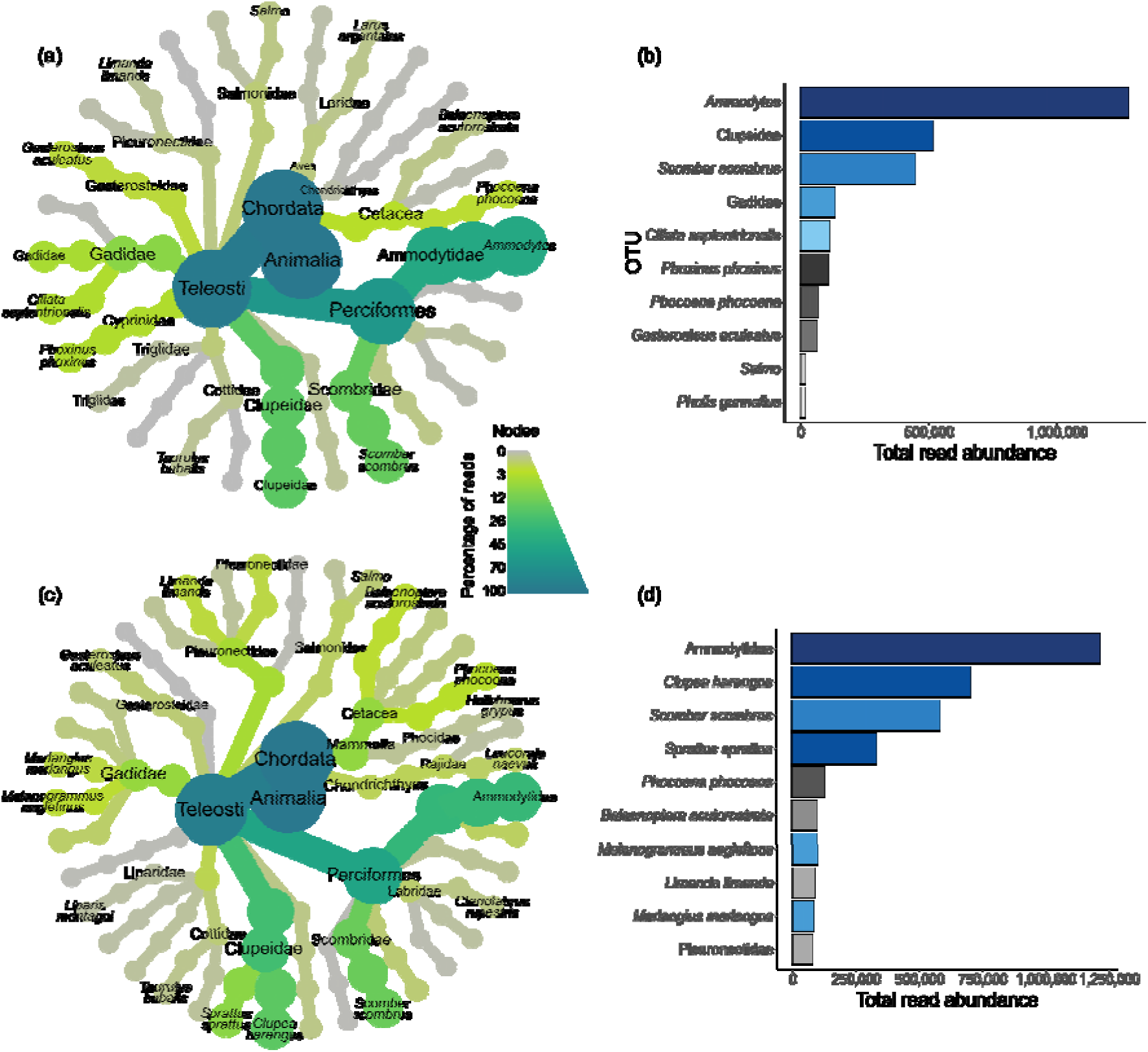
Heat trees showing the detected taxa with more than 2000 reads for (a) the MarVer1 primer set, and taxa with more than 3000 reads for (c) the MarVer3 primer set. The size and colour of the nodes represents the proportion of reads that a taxa contributes too. Bar charts displaying the ten OTUs with the most abundant read counts for (b) MarVer1, and (d) MarVer3. The colour highlights OTUs belonging to the same family.

**Table 1.** Comparison of the taxa composition across the vertebrate primer markers MarVer1 and MarVer3.

| Class | MarVer1<br>Total reads<br>(% of reads) | MarVer1<br>OTUs detected | MarVer3<br>Total reads<br>(% of reads) | MarVer3<br>OTUs detected |
| --- | --- | --- | --- | --- |
| All | 2,880,775 | 5 | 4,013,997 | 80 |
| Teleosts | 2,763,655 (96 %) | 41 | 3,653,206 (91 %) | 63 |
| Mammals | 89,835 (3 %) | 5 | 294,778 (7 %) | 6 |
| Chondrichthyes | 5726 (<1 %) | 3 | 62,980 (1.5 %) | 5 |
| Birds | 21,113 (<1 %) | 6 | 3033 (<1 %) | 6 |
| Cephalaspidomorphi<br>(lamprey) | 447 (<1%) | 1 | 0 | 0 |

The top three most abundant OTUs across both markers were forage fish species, with sandeels having the highest read counts, accounting for 30% and 44% of the reads for MarVer1 and MarVer3 respectively (Figure 3). This was followed by the Clupeidae family, which could only be resolved at species level to herring and sprat with MarVer3, and then mackerel (*Scomber scombrus*). The Gadidae family had the fourth highest reads for MarVer1, whilst species from the Gadidae family, haddock (*Melanogrammus aeglefinus*) and whiting (*Merlangius merlangus*) had seventh and ninth highest read counts respectively for MarVer3. Two cetacean species, minke whales and harbour porpoises, appeared in the ten most abundant reads for MarVer3, but only the harbour porpoise was present among the ten most abundant taxa for MarVer1, although minke whales had the eleventh most abundant reads.

### Composition of broader eukaryotic taxa detected

We retrieved 1,769,650 reads in total with our 18S primer set, belonging to 122 different eukaryotic families (Table 2). These sequences largely comprised families belonging to either the Animalia (48% total reads) or Chromista (41% total reads) kingdoms (Figure 4a). Reads in the Animalia kingdom were dominated by Calanidae and Acartiidae copepod families from the Maxillopoda class, which together accounted for 42% of the total reads (Figure 4). The most abundant families, Leptocylindraceae and Calciodinellaceae, from the diatom (Bacillariophyceae) and dinoflagellate (Dinophyceae) classes respectively, accounted for 18% of the total reads. The next most abundant classes were haptophytes (Prymnesiophyceae), also from the Chromista kingdom, and fungi (Phycomycota). Didiniidae and Strombidiidae belong to the ciliates class, but Ciliata was removed as potential contamination when analysed at class level due to other ciliate families being abundant in field blanks.

**Figure 4.**
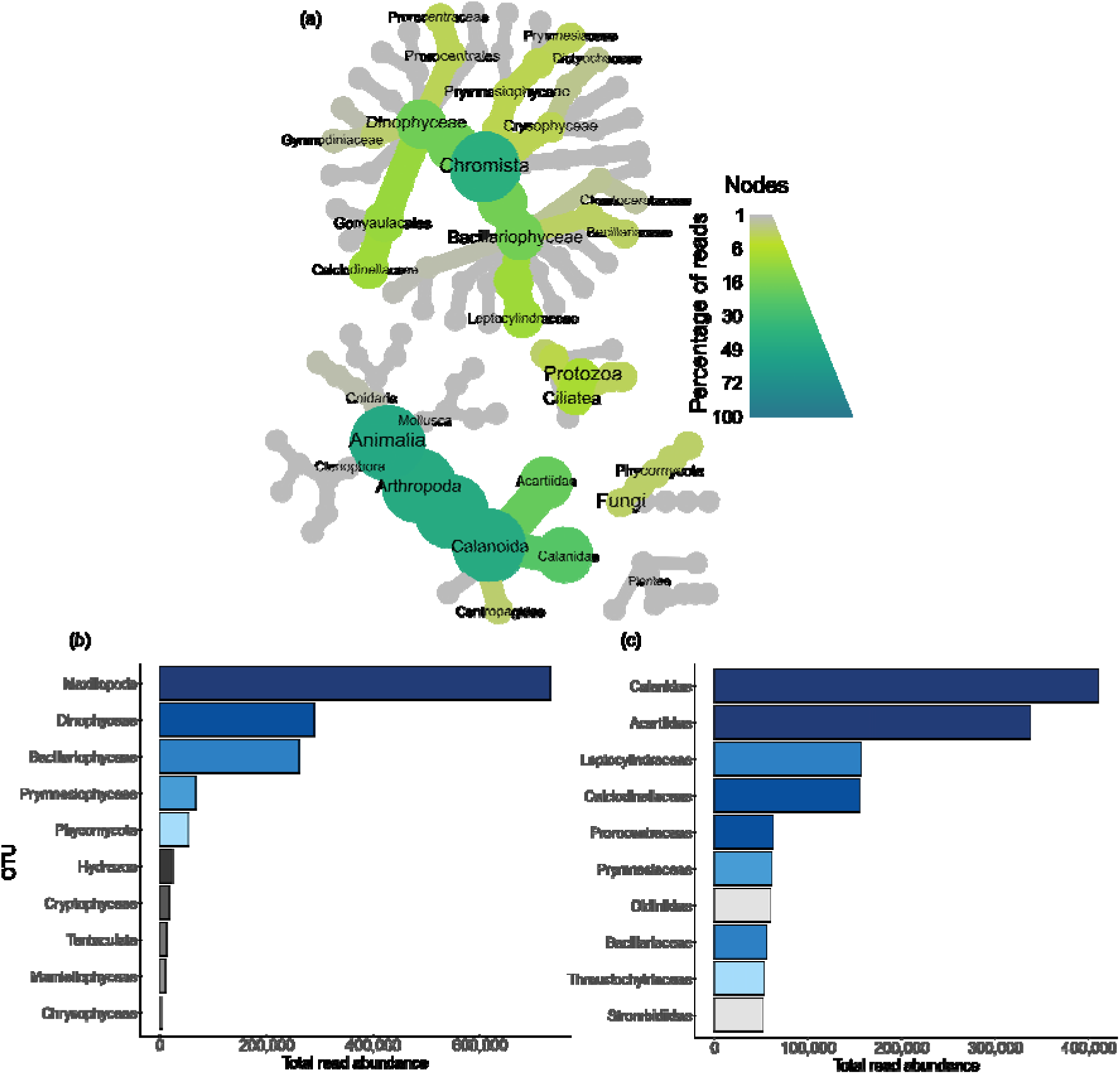
(a) Heat tree showing taxa detected with more than 500 reads by the 18S primer set. The size and colour of the node represents the proportion of reads contributed by the taxa. Bar charts with the ten most abundant OTUs for 18S at (b) Class taxonomic groups, and (c) Family taxonomic groups. Colours indicate which class the families belong in. Didiniidae and Strombidiidae families belong to the Ciliatea class which is removed as contamination at class level.

**Table 2.** Composition of sequencing reads within different taxonomic kingdoms retrieved from the V9 region of 18S rRNA, and the number of OTUs detected at family level.

| Kingdom | 18S - Total Reads<br>(% of reads) | 18S – OTUs<br>detected |
| --- | --- | --- |
| All | 1,769,650 | 122 |
| Animalia | 845,096 (48%) | 47 |
| Chromista | 727,542 (41%) | 48 |
| Protozoa | 130,151 (7%) | 14 |
| Fungi | 53,767 (3 %) | 5 |
| Plantae | 13,092 (<1%) | 8 |

### Temporal trends in community composition

Sandeels (Ammodytidae) had higher proportions of read counts between June and August with MarVer1 and June and July with MarVer3 (Figure 5). With the ensemble OTU index, sandeels were an indicator OTU for June, July and August (specificity = 0.98, sensitivity = 0.82, stat = 0.9, p = 0.005) (Appendix Table A5). Minke whales (Balaenopteridae) were most prevalent in June and July, and an indicator for these months (specificity = 0.93, sensitivity = 0.88, stat = 0.91, p = 0.02), whilst harbour porpoises (Phocoenidae) had higher read proportions between June and August and were an indicator for these months (specificity = 0.9999, sensitivity = 0.87, stat = 0.93, p = 0.005). Mackerel (Scombridae) were detected across our full temporal scale but made up a greater proportion of reads between August and October. Similarly, herring and sprat (Clupeidae) were also detected across all months but had the highest read proportions in June and September/October.

**Figure 5.**
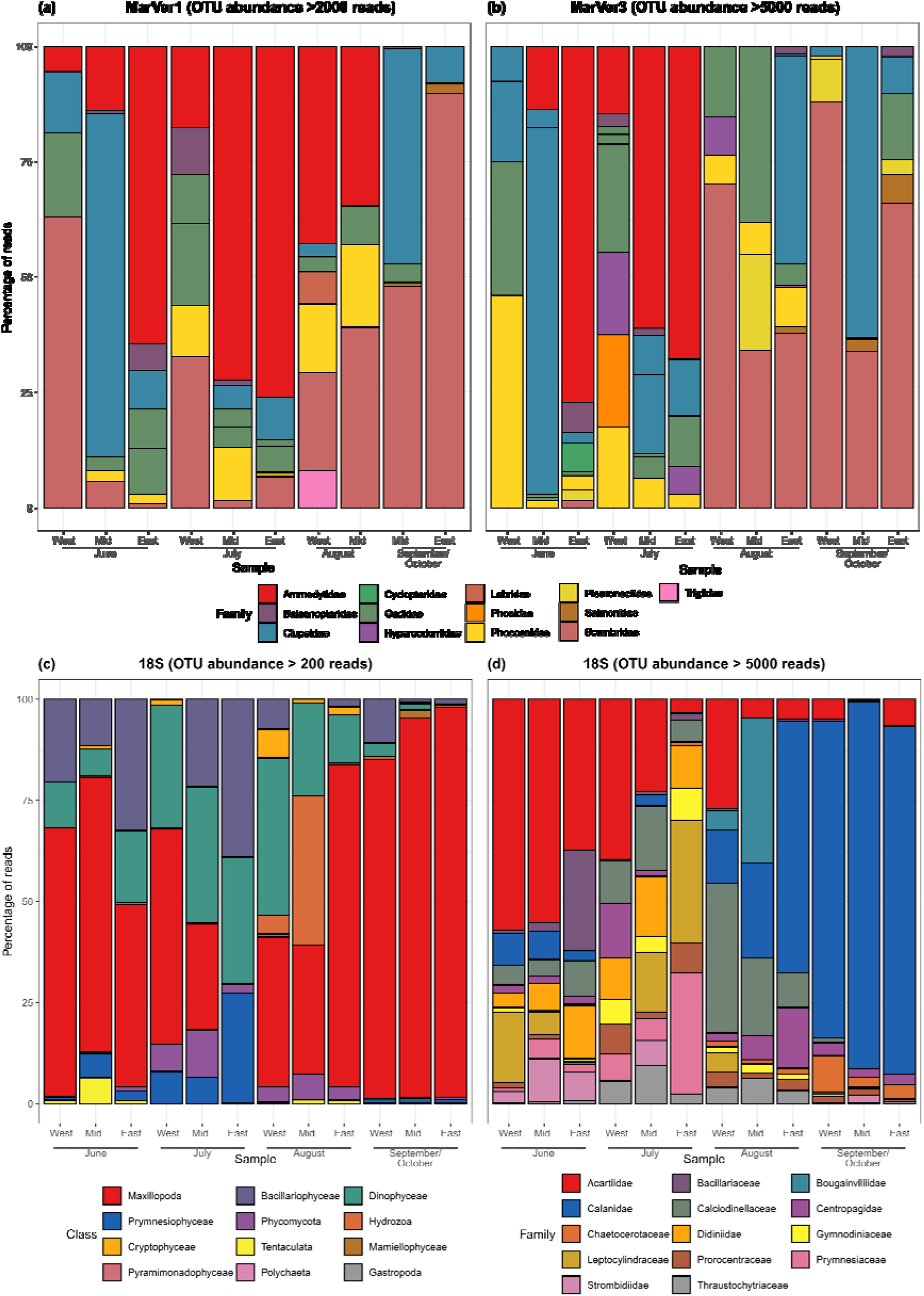
Stacked bar charts showing the most abundant vertebrate families for the primer sets (a) MarVer1 and (b) MarVer3, (c) the most abundant 18S classes and (d) the most abundant 18S families from the twelve fixed sampling points. Two fixed samples are missing for MarVer1 due to failed amplification.

Maxillopoda made up a greater proportion of reads in June and September/October, with Acartiidae being the most prominent copepod family in early sampling months and Calanidae in the latter months (Figure 5). Calanidae were an indicator for August and September/October (specificity=0.93, sensitivity=0.78, stat=0.85, p=0.001) (Appendix Table A6). Diatoms (Bacillariophyceae) were most abundant in June and July, predominantly represented by Leptocylindraceae and Bacillariaceae families, shifting to the Chaetocerotaceae family in the latter sampling months. Dinoflagellates were most prevalent in July and August, with the Calciodinellaceae family contributing the greatest proportion and being an indicator species for these months (specificity=0.87, sensitivity=1, stat=0.93, p=0.001). Ciliate families, Didiniidae and Strombidiidae, were more abundant in June and July, and Didiniidae (specificity=0.98, sensitivity=1, stat=0.989, p=0.001) was an indicator for these months, while Strombidiidae was an indicator for June, July and September/October (specificity=0.996, sensitivity=0.93, stat=0.964, p=0.001). The haptophyte family Prymnesiaceae was also most abundant in June and July, and an indicator species for June, July and August (specificity=0.998, sensitivity=1, stat=0.999, p=0.001). Hydrozoa were most frequently detected in August, represented by the Bougainvilliidae family, which was also an indicator OTU for this month (specificity=0.98, sensitivity=0.58, stat=0.76, p=0.001).

Alpha diversity, with the Shannon Index, did not significantly differ between sampling months for our ensemble vertebrate OTU index or our eukaryote OTU index, although all primer sets had lowest alpha diversity in September/October (Figure 6a). PERMANOVA shows that community composition differs between sampling months for both our vertebrate OTU index (*adonis*; *df* = 3, *F* = 2.35, *R^2^* = 0.12, *p* = 0.001) and our eukaryote OTU index (-*adonis*; *df* = 3, *F* = 9.6, *R^2^* = 0.35, *p* = 0.001). For eukaryotes, community composition was significantly different between all months (pairwise adonis, P < 0.01), whereas for vertebrate communities, the communities in June and July differed from the communities in August and September/October (pairwise adonis, P < 0.05). NMDS analysis also revealed distinct communities per month for both vertebrates and eukaryotes (Figure 6).

**Figure 6.**
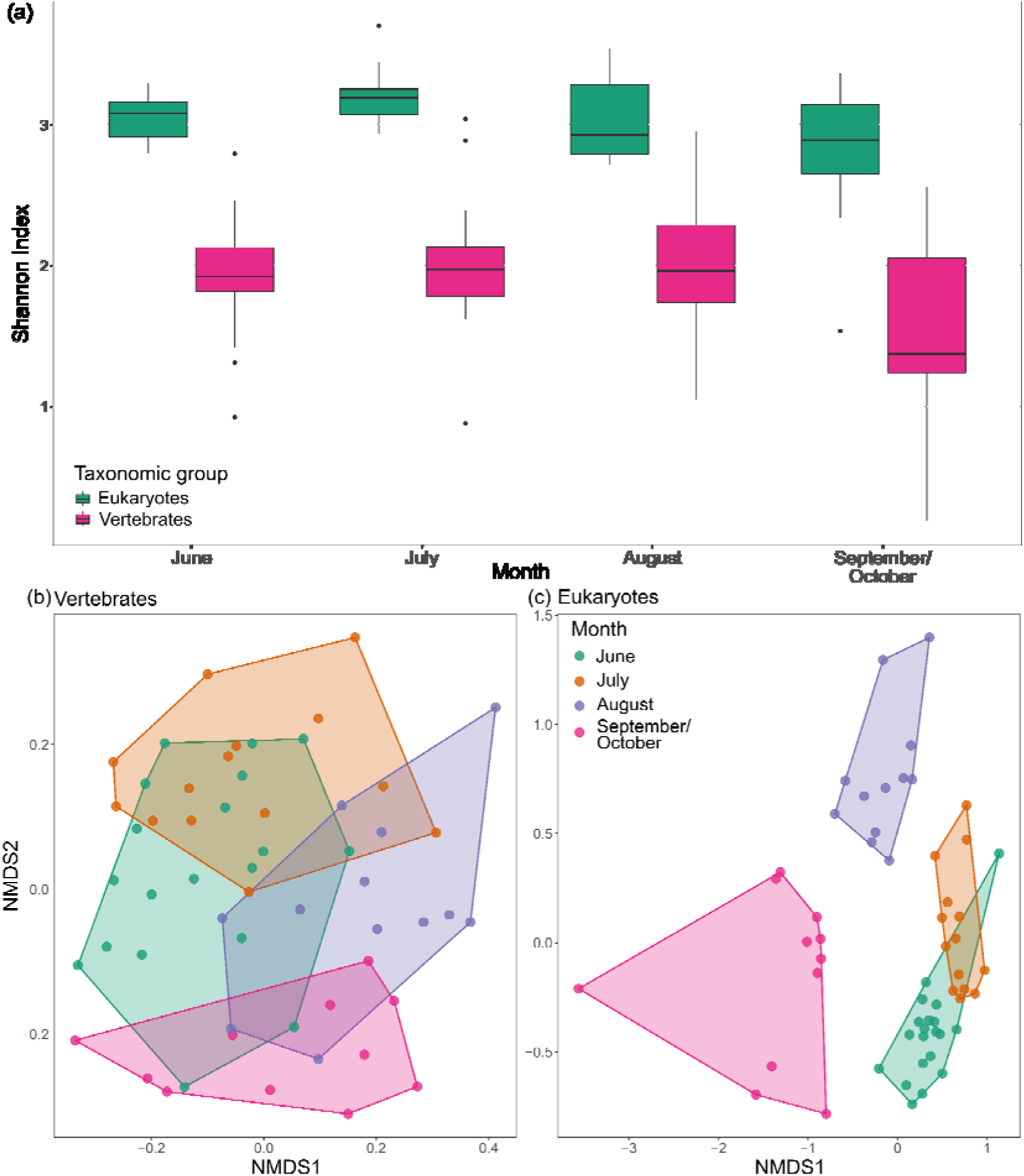
(a) Box plots showing alpha diversity for eukaryotes and vertebrates across different sampling months, and NMDS plots for (b) our ensemble vertebrate OTU index (k = 2, stress = 0.262) and (c) eukaryotic OTU index (k = 2, stress = 0.204), as partitioned by sampling month.

### Spatial trends in community composition

We detected spatial partitioning between cetaceans commonly found in the Moray Firth, with bottlenose dolphins occurring in greatest abundance closest to shore, and with minke whales and harbour porpoises being more abundant in samples greater than 2.5 km from shore (Figure 7a). Bottlenose dolphins were an indicator for the near and middle near categories (specificity = 0.996, sensitivity = 0.82, stat = 0.91, p = 0.005), whilst the harbour porpoise was an indicator for the middle near, middle far and far categories (specificity = 0.98, sensitivity = 0.84, stat = 0.9, p = 0.005). The availability and abundance of different cetacean prey species also varied across distance from shore (Figure 7b). Sandeels were found in similar abundance across all depths, whilst clupeids were most prominent between 7 and 10 km from the coast. Salmonids were most abundant within 1.2 km of the shore, whilst the Gadidae family were most abundant in the near and middle near categories as well as the far category, probably represented by different species within the Gadidae family.

**Figure 7.**
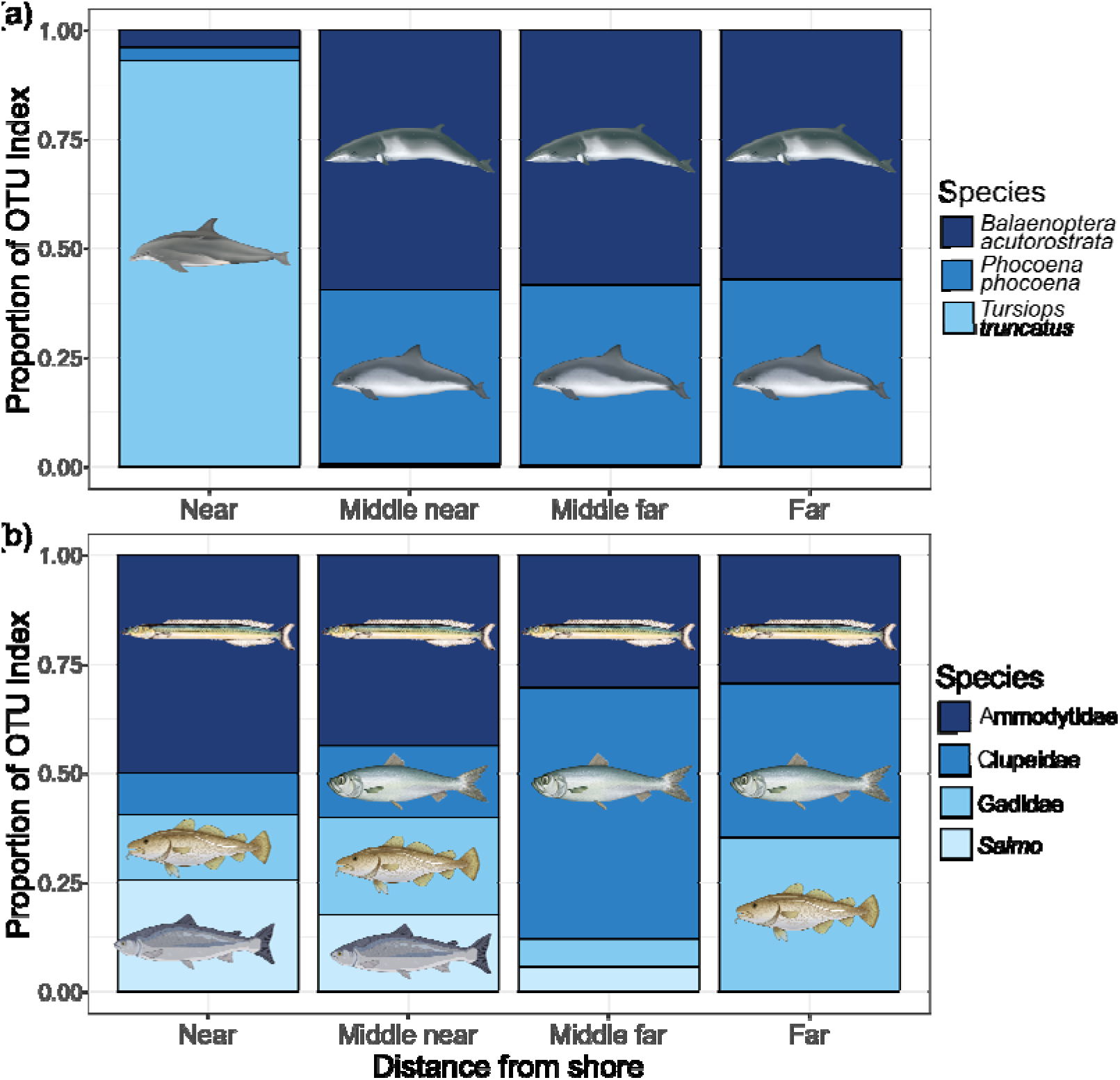
Bar charts showing the proportion of the OTU index for (a) common cetacean species in the study area: minke whales (*Balaenoptera acutorostrata*), harbour porpoises (*Phocoena phocoena*) and bottlenose dolphins (*Tursiops truncatus*); and (b) their dominant prey species across different distances from shore: near <1.2 km, middle near between 2.5 and 7 km, middle far between 7 and 10 km, and far > 10 km (up to 16.1 km) offshore.

Alpha diversity, described with the Shannon Index, significantly differed with distance from shore for both vertebrates (Kruskal–Wallis χ^2^ = 15.68, df = 3, p <0.001) and eukaryotes (Kruskal–Wallis χ^2^ = 18.12, df = 3, p <0.001). In both cases, the nearshore community had significantly higher alpha diversity compared to all groups further offshore (Pairwise Wilcoxon Rank Sum test <0.05) (Figure 8a). PERMANOVA analyses found beta diversity to significantly differ with distance from shore for both vertebrate (*adonis*: *df* = 3, *F* = 2.03, *R^2^* = 0.10484, *p* = 0.001) and eukaryote communities (*adonis*: *df* = 3, *F* = 1.8274, *R^2^* = 0.09, *p* = *0.002*), although within-community variance was not homogenous (*F* = 2.9, p<0.05) for eukaryotes. The vertebrate nearshore community differed significantly from all offshore categories, whilst the eukaryotic nearshore community differed from the middle-far and far categories (pairwise adonis, p < 0.05). NMDS showed that the vertebrate nearshore community was distinct from communities further offshore but displayed great overlap between distance from shore categories for eukaryotes (Figure 8). The nearshore community had the highest number of OTU indicators for both eukaryotic and vertebrate communities (Figure 9). 20 eukaryotic OTUs were identified as indicators for the nearshore community, including 6 families that were only found in nearshore samples; red (Rhodomelaceae) and brown (Dictyotaceae) algae families, calcareous sponges (Leucosoleniidae), tunicates (Molgulidae), bryozoans (Membraniporidae) and cyclopoid copepods (Archinotodelphyidae) (Appendix Table A7). Ten vertebrate OTUs were recognised as indicators, most of which were species commonly residing in shallow depths less than 50 metres, such as the rock gunnel, ballan wrasse (*Labrus bergylta*), eelpout (*Zoarces*), Montagu’s seasnail (*Liparis montagui*), painted goby (*Pomatoschistus pictus*) and the leopard spotted goby (*Thorogobius ephippiatus*) (Appendix A8).

**Figure 8.**
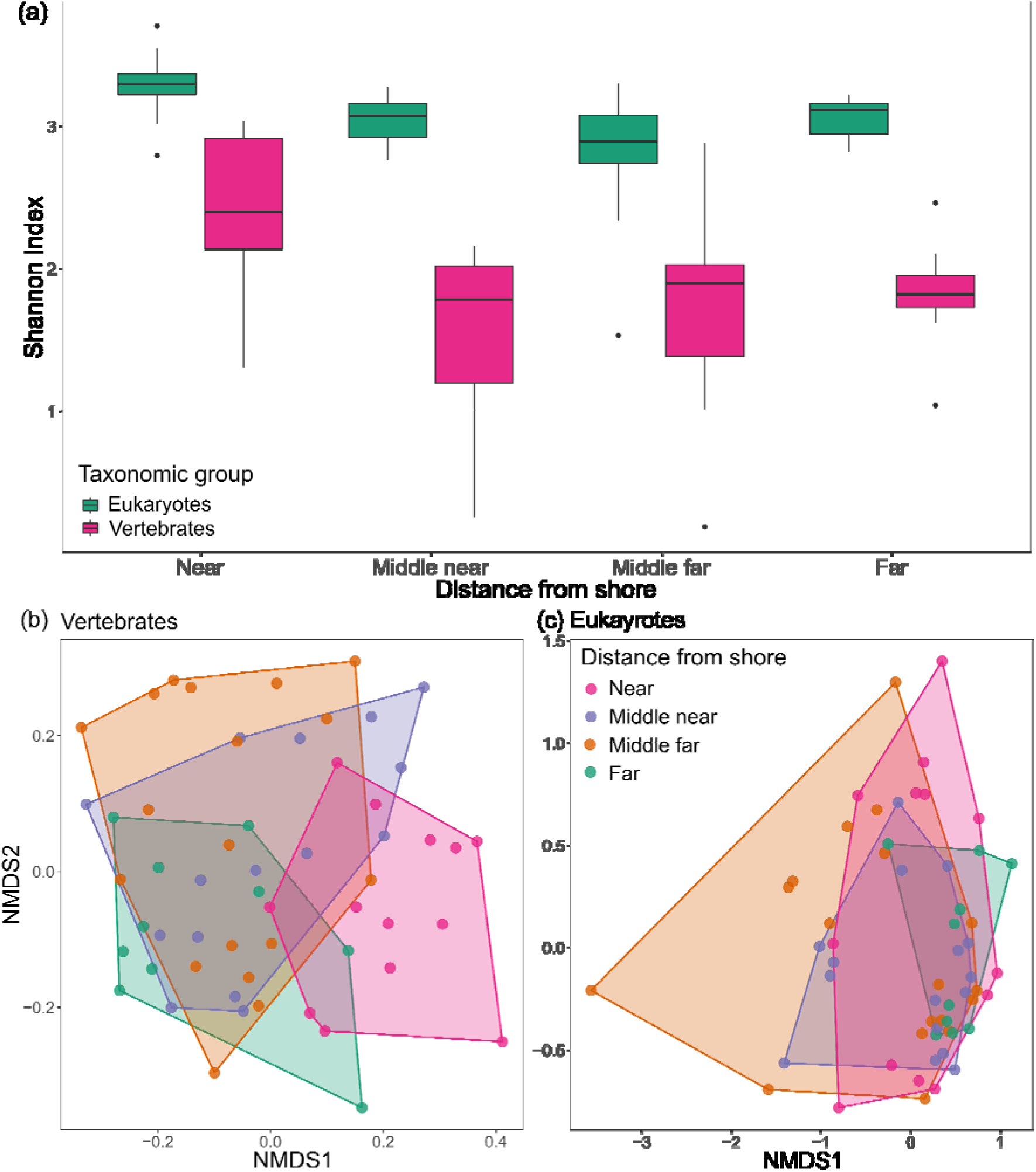
(a) Box plots showing alpha diversity for eukaryotes and vertebrates across different distances from shore, and NMDS plots for the (b) ensemble vertebrate OTU index (k = 2, stress = 0.2) and the (c) eukaryotic OTU index (k = 2, stress = 0.18) partitioned by distance from shore.

**Figure 9.**
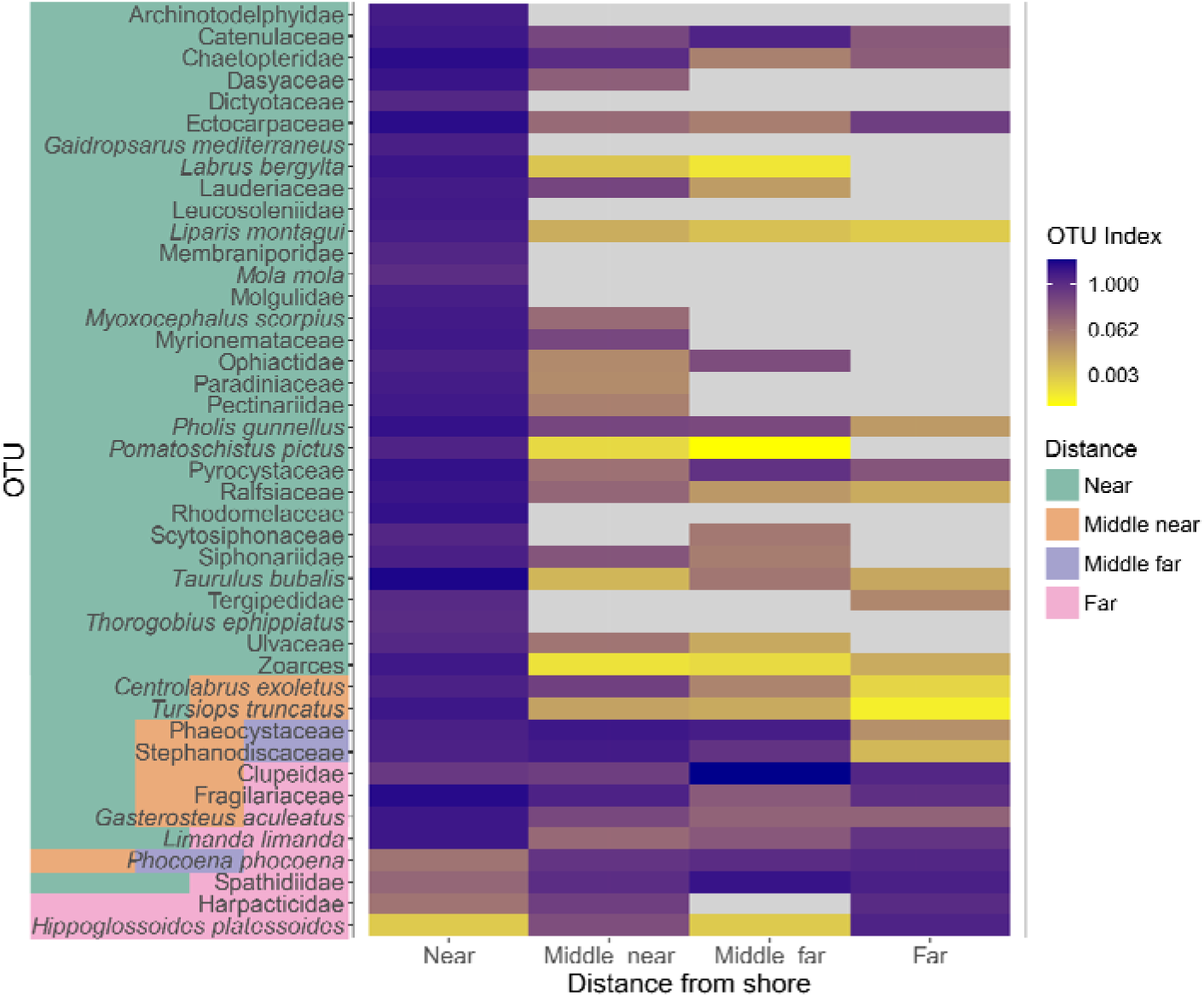
Heat map showing indicator OTUs for distance from shore categories across vertebrate and broader eukaryotes. OTUs are coloured to show which distance from shore category or combination of categories they are an indicator species for.

### Environmental drivers of community composition

MRM revealed that both temporal, *i.e.,* distance between days, and spatial drivers, *i.e.*, geographical distance, are positively correlated with vertebrate beta diversity (Table 3). For eukaryotic beta diversity, difference between days was also positively correlated, along with sea surface temperature. For both vertebrates (coeff = 0.32) and eukaryotes (coeff = 0.8), difference between days had the greatest influence on beta diversity.

**Table 3.** Results of minimal multiple regression distance models for eukaryote and vertebrate OTU indexes.

| Eukaryote minimal MRM model |  |  | Vertebrate minimal MRM model |  |  |
| --- | --- | --- | --- | --- | --- |
| Variable | Coefficient | P value | Variable | Coefficient | P value |
| Intercept | 68.16 | 1.00 | Intercept | 443.15 | 1.00 |
| Sea Surface Temperature | 0.08 | 0.02 | Geographic distance | 0.1 | 0.02 |
| Difference in calendar days | 0.8 | 0.0001 | Difference in calendar days | 0.32 | 0.0001 |
| Full model statistics |  |  |  |  |  |
| $R^2 = 0.66$ | | | $R^2 = 0.12$ | | |
| P value = 0.0001 |  |  | P value = 0.0001 |  |  |

## Discussion

Limited knowledge of prey availability and distributions frequently prevents us from fully understanding heterogeneity and seasonality in marine mammal distributions (Pendleton et al., 2020; Szesciorka et al., 2023). Given that MPAs for marine mammals commonly focus on foraging grounds, it is important to understand their seasonal habitat use within these areas and predict the long-term stability or locational shifts in populations (Notarbartolo Di Sciara et al., 2016). Here, we discovered that the availability and abundance of key cetacean forage fish species varied both seasonally and with distance from shore, providing important insights into the habitat use of the marine mammal species present in a newly implemented MPA in the southern Moray Firth, north-east Scotland. Further, we recovered spatiotemporal trends in overall community composition which were mirrored in both vertebrate and broader eukaryotic communities, indicating a highly connected ecosystem. The North Sea is warming faster than global averages (Holt et al., 2012), and forage fish present in the basin are reliant on specific substrates making them particularly vulnerable to warming temperatures as they have limited options to migrate further north to suitable habitat patches (Frederiksen et al., 2011; Petitgas et al., 2013). Therefore, continued monitoring will be essential to track prey availability for marine mammals, and to detect asynchronous timings in predator-prey presence which could have negative cascading effects throughout the ecosystem (Silber et al., 2017).

### Seasonality in community composition

Temporal drivers had the strongest effects on both eukaryotic and vertebrate communities (Table 2.3), despite sampling having been undertaken over a relatively short time scale (June to October). Communities in June and July were more similar compared to communities in the latter sampling months, reflecting in shifts in the most abundant OTUs over time (Figure 5). For example, Acartiidae was the most prevalent copepod family in June and July but this switched to Calanidae from August onwards. Similarly, sandeels were the most abundant vertebrate in June and July, but from August onwards were replaced by mackerel as the most commonly detected taxon. The similarity in temporal patterns between eukaryotic and vertebrate communities could be indicative of strong connectivity between the taxonomic groups and tight coupling of interactions. Previous declines in the North Sea zooplankton biomass due to warming temperatures, have been linked to declines in forage fish biomass, and failure of forage fish populations to recover after enforcement of stricter fishing regulations (Clausen et al., 2018; Lindegren et al., 2018). Conversely, forage fish can also exert top-down control on zooplankton (Fauchald et al., 2011; Lynam et al., 2017). Given that SST is a driver of our eukaryotic community (Table 3), and the North Sea is warming three times more quickly than the global average, our results suggest future changes in zooplankton composition (Belkin, 2009; Emeis et al., 2015), can be expected. Whole ecosystem-based monitoring incorporating eDNA tools will be essential to detect early changes and track prospective cascading effects throughout the ecosystem.

### Community composition changes with distance from shore

The nearshore community, within 1.2 km of the shore and less than 25 metres depth, had higher species richness and significantly differed from those communities sampled further offshore. This observation could partially be a methodological artefact resulting from the 4m sampling depth that could increase the likelihood of detecting benthic species in shallower water (Figure 8). However, OTUs detected in higher abundance or only in nearshore samples represented species and families that are typically constrained to shallower depths. For example, two algae families, Dictyotaceae and Rhodomelaceae, were only found in nearshore samples which corresponded to the depth limits of algal growth in the North Sea (Pehlke & Bartsch, 2008; van der Stap et al., 2016). Similarly, American plaice (*Hippoglossoides platessoides*), a demersal species, was most prevalent in those samples collected furthest from shore, between 80 and 100 metres depth, suggesting strong mixing of eDNA within the water column, or detections of their pelagic larval phase (Walsh, 1994). Clear spatial partitioning between common cetacean species was detected with eDNA, matching distributions inferred from visual surveys (Robinson et al., 2007). For example, bottlenose dolphin eDNA was far more abundant in the nearshore samples, with long term visual surveys showing this population occupies depths less than 25 metres (Culloch & Robinson, 2008). Both minke whales and harbour porpoises were detected across all distances from shore, but in greater abundance offshore (>2.5 km), also corroborating long term visual survey efforts (Robinson et al., 2007). In addition, minke whale sighting samples had significantly higher abundance compared to control samples (Appendix A1). Transport of eDNA from its source by currents and tides is an ongoing concern for the incorporation of eDNA into monitoring of the marine environment (Andruszkiewicz et al., 2019; Hansen et al., 2018). Our results contribute to a growing body of work demonstrating that eDNA provides a relatively local signal from species in the marine water column, suggesting that eDNA degrades rapidly or becomes diluted beyond detectable limits quickly as it is transported away from its source (Hansen et al., 2018).

### Minke whale habitat use in relation to prey species

Minke whales are the only species included as a biodiversity feature in the Moray Firth Southern Trench MPA, as a result of the area being an important foraging ground attracting above average abundances of minke whales (Robinson et al., 2009). Current lack of knowledge on the spatiotemporal availability of targeted prey species has hampered identification of important focal areas within the MPA for minke whales. The dominant minke whale prey species, such as sandeels and clupeids, were the most abundant vertebrate species detected with our primer sets (Figure 3). Abundances vary throughout the foraging season which would account for the dietary plasticity exhibited by these whales (Robinson et al., 2023). Sandeels were most abundant during June and July, when they are foraging in the water column, but declined from August onwards, when they return to burrows in the sediment (Henriksen et al., 2021) (Figure 2.5). Meanwhile, clupeids were most abundant in June and September. The early peak in abundance coincides with the main spawning period of sprat, whilst the latter peak likely represents the arrival of juvenile sprat and herring to overwinter in the firth (Thompson et al., 1991). Juvenile minke whales target yearling sandeels throughout the foraging season, whilst adults target larger sandeels before switching to sprat and juvenile herring as they become more abundant (Robinson et al., 2023). Juvenile minkes are also found at shallower depths, while adults prefer deeper, offshore waters, corresponding with the distance from shore that their targeted prey species are found (Robinson et al., 2023). Accordingly, while sandeels were detected across all depths, clupeids were detected in greatest abundance between 7 and 10 km offshore, at depths between 50 and 120 metres (Figure 7). Both sandeels and clupeids are reliant on specific bottom substrates, making them vulnerable to climate-induced depletions as they are restricted in their ability to find new habitat patches that would facilitate northward migration (Frederiksen et al., 2011; Petitgas et al., 2013). Marine mammals generally respond to prey depletions by switching to alternative prey species or moving to new foraging grounds (Agardy et al., 2019). Elsewhere, mackerel and gadoids, such as cod, haddock and whiting, have become more important components of minke whale diets as their preferred prey, such as krill species and capelin (*Mallotus villosus*), have declined (Víkingsson et al., 2014; Windsland et al., 2007). We detected high abundances of mackerel and gadoids, suggesting that potential alternative prey sources exist in the Moray Firth that could support similar climate-induced prey switches. However, it should be noted that alternative species have lower energy value densities compared with sandeels (Ransijn et al., 2019; Van Pelt et al., 1997), with potential implications for population demography (Österblom et al., 2008; Spitz et al., 2012).

### Species of conservation interest

eDNA also detected other species of conservation interest within the Southern Trench MPA. For example, the detections of Sowerby’s beaked whales were unexpected, as no definitive live sightings have been recorded in the North Sea, despite high survey efforts, and strandings are rare (MacLeod et al., 2004). In view of the long periods spent beneath the surface, beaked whales are notoriously difficult to detect visually, so eDNA could be an important tool to improve our understanding of this species distribution and conservation status, given that they are listed as data deficient in the IUCN ‘Red List’ (Hooker et al., 2019). Our eDNA detections were coincident with a Sowerby’s beaked whale stranding, with at least one sample preceding the event, thus the eDNA and stranding are indicative of the species presence in the area.

The North Sea Atlantic bluefin tuna fishery collapsed in the 1960s and records have been sparse ever since, although in recent years, sporadic observations suggest that they are making a return to UK waters (Horton et al., 2021). Bluefin tuna were detected in seven samples collected in June, providing an important record of bluefin tuna returning to historical foraging grounds or potentially expanding their migration routes in between foraging and overwintering/spawning grounds (Horton et al., 2021).

We also detected the critically endangered European eel, for which the timing of migration and movement patterns are currently poorly understood around Scotland (Malcolm et al., 2010). Detections of European eel peaked in August, which could be related to either adult eels leaving rivers to return to their breeding grounds in the Sargasso sea, or the arrival of juvenile glass eels (Malcolm et al., 2010). The invasive pink salmon was detected across all sampling months but occurred in greatest abundance in nearshore samples. Pink salmon have been found in low abundance in Scottish rivers for over 50 years, particularly in the River Spey which flows into the Moray Firth at the western boundary of the study area (Armstrong et al., 2018). It is speculated that pink salmon fry enter the sea at the onset of winter, leading to low survival rates, but there is currently no evidence to support this (Armstrong et al., 2018). Accordingly, eDNA could provide an additional tool to monitor temporal dynamics of pink salmon in Scottish rivers in relation to the development of management strategies (Gargan et al., 2022).

### Limitations and future work

One of the biggest limitations of using eDNA metabarcoding to explore predator-prey dynamics is the inability to distinguish between different age classes (Hansen et al., 2018), as predators often preferentially target prey of certain sizes (Robinson et al., 2023; Visser et al., 2021). In particular, spawning events have been observed to increase the abundance of read counts retrieved in eDNA studies, but larval forms are unlikely to be suitable prey for piscivorous marine mammals (Di Muri et al., 2022; Ratcliffe et al., 2021). However, eDNA can provide information about where and when to carry out more intensive surveys to retrieve parameters such as the age-class structure of fishes present. Similarly, abundance estimates of cetaceans are important to monitor trends in population sizes (Hammond et al., 2021), but we are unable to conclude from eDNA how many individual cetaceans were using the MPA and whether there was turnover of individuals across the season. We only collected samples during one foraging season, but the number of minke whales visiting the area is known to vary interannually (Robinson et al., 2009), so it would be interesting to investigate broader interannual community changes and relate these to the number of minkes and environmental drivers. Changeable weather conditions further limited where samples could be collected, with less samples collected offshore from August onwards, and with more westerly samples collected in July compared to easterly samples in September/October (Figure 2). These spatial differences could therefore have influenced the temporal signals retrieved, although samples collected <2.5 km offshore were all very similar in their community composition, so we believe this had minimal impact on the temporal trends observed. Future work could be targeted to extend the sampling area outside of the MPA in order to evaluate the informativeness of eDNA for tracking the effectiveness of management actions within the MPA (Dunham et al., 2020). Extending eDNA monitoring to other UK MPAs designated for cetaceans, and to areas subject to current and future industrial activity, e.g. offshore windfarms (Isaksson et al., 2023), would be informative for comparing and contrasting their respective community compositions, as well as building the foundation of long term eDNA datasets that could contribute to tracking ecosystem changes due to climate warming.

## Conclusion

Our results demonstrate that eDNA metabarcoding can serve as a powerful tool to monitor marine mammals and their prey species simultaneously, improving the understanding of marine mammal habitat use on their foraging grounds. eDNA approaches could support monitoring of MPAs focused on these foraging areas by informing us about seasonal distribution changes and heterogenous habitat use, as well as contributing to long-term dynamic management of foraging areas as prey species shift their distribution in response to global warming (Notarbartolo Di Sciara et al., 2016). To this end, we provide an important baseline characterisation of community composition within the Southern Trench MPA in the outer Moray Firth against which future changes can be evaluated. Key forage fish species, such as sandeels and clupeids, account for 86% of the total fish biomass in the Moray Firth and are the main prey species for many piscivorous fishes, seabirds and marine mammals alike (Greenstreet et al., 1998). These species are especially vulnerable to global warming, and thus monitoring will be essential to inform potential changes in abundance and to assess how predators will respond (Frederiksen et al., 2011; Petitgas et al., 2013).

## Data Accessibility

The data will be archived in the NCBI BioProject repository at the time of publication.

