## Appendix for "Environmental DNA reveals fine scale spatial and temporal variation of prey species for marine mammals in a Scottish marine protected area"

| Appendix Table A1 | Metadata for each eDNA sampling point. |
| --- | --- |
| Appendix Table A2 | List of taxa detected by the different primer sets, MarVer1, MarVer3 and 18S. |
| Appendix Table A3 | Foreign OTUs detected with MarVer1 were compared to native relatives from the same genus or family. Reads were either reassigned to the native relative, removed due to low abundance or identified as a potential invasive. |
| Appendix Table A4 | Foreign OTUs detected with MarVer3 were compared to native relatives from the same genus or family. Reads were either reassigned to the native relative, removed due to low abundance or identified as a potential invasive. |
| Appendix Figure A1. | OTU index for minke whales in control versus sighting samples for (a) MarVer1 and (b) MarVer3. |
| Appendix Table A5 | Indicator species analysis for vertebrates per trip. |
| Appendix Table A6 | Indicator species analysis for eukaryotes per trip. |
| Appendix Table A7 | Indicator species analysis for eukaryotes for distance from shore categories. |
| Appendix Table A8 | Indicator species analysis for vertebrates for distance from shore categories. |

Appendix Table A1. Metadata for each eDNA sampling point. Samples were named such that the first letter corresponded to the type of sample (C = Control, F = Fixed, S = Sighting), the first number to the monthly trip (1 = June, 2 = July, 3 = August, 4 = September/October) and the second number to the sample site (1 to 3 for fixed samples from west to east, or the chronological order of samples collected for that trip for control and sighing samples.)

| **Name** | **Date** | **Time** | **Longitude** | **Latitude** | **Bathymetry (m)** | **Sea surface temperature (ºC)** | **Chlorophyll *a* (mg/m^3^)** | **Distance from shore (m)** | **Distance from shore category** |
| --- | --- | --- | --- | --- | --- | --- | --- | --- | --- |
| C1.1 | 23/06/2021 | 16:17 | -2.38349 | 57.77308 | -134 | 13.4 | 0.8069 | 9,840 | Middle far |
| C1.2 | 24/06/2021 | 10:47 | -2.55324 | 57.72608 | -72 | 13.55 | 0.5582 | 4,896 | Middle near |
| C1.3 | 24/06/2021 | 12:14 | -2.3384 | 57.7856 | -99 | 13.81 | 0.8025 | 10,188 | Far |
| C1.4 | 24/06/2021 | 12:39 | -2.32011 | 57.73321 | -70 | 13.12 | 1.2222 | 4,278 | Middle near |
| C1.5 | 27/06/2021 | 13:22 | -2.28656 | 57.69763 | -20 | NA | 1.63 | 352 | Near |
| C1.6 | 30/06/2021 | 12:23 | -2.7032 | 57.69119 | -7 | NA | NA | 393 | Near |
| C1.7 | 30/06/2021 | 14:37 | -2.84249 | 57.73908 | -27 | 14.21 | 1.2632 | 3,535 | Middle near |
| C1.8 | 30/06/2021 | 16:50 | -2.57757 | 57.68753 | -21 | 13.79 | 2.5682 | 458 | Near |
| F1.1 | 30/06/2021 | 14:55 | -2.82517 | 57.76298 | -33 | 14.28 | 1.0743 | 6,333 | Middle near |
| F1.2 | 23/06/2021 | 13:48 | -2.54138 | 57.76318 | -104 | 13.46 | 0.6049 | 9,065 | Middle far |
| F1.3 | 27/06/2021 | 15:33 | -2.26 | 57.7637 | -97 | 13.01 | 1.2358 | 7,765 | Middle far |
| S1.1 | 23/06/2021 | 12:56 | -2.54576 | 57.74366 | -91 | 13.36 | 0.5796 | 6,889 | Middle near |
| S1.2 | 23/06/2021 | 14:25 | -2.51157 | 57.82474 | -85 | 13.74 | 0.6064 | 16,144 | Far |
| S1.3 | 23/06/2021 | 15:25 | -2.38127 | 57.83258 | -91 | 13.56 | 0.8228 | 15,928 | Far |
| S1.4 | 24/06/2021 | 11:25 | -2.49849 | 57.79097 | -77 | 14.07 | 0.6134 | 12,755 | Far |
| S1.5 | 24/06/2021 | 11:48 | -2.43766 | 57.79806 | -90 | 14.07 | 0.6742 | 13,794 | Far |
| S1.6 | 27/06/2021 | 15:21 | -2.21621 | 57.76389 | -91 | 12.25 | 1.5723 | 8,837 | Middle far |
| S1.7 | 28/06/2021 | 12:12 | -2.55214 | 57.74915 | -87 | 13.36 | 0.9526 | 7,407 | Middle far |
| S1.8 | 28/06/2021 | 15:59 | -2.50392 | 57.78123 | -82 | 12.94 | 1.148 | 11,625 | Far |
| S1.9 | 30/06/2021 | 15:18 | -2.69178 | 57.76373 | -69 | 13.96 | 1.4653 | 8,023 | Middle far |
| C2.1 | 14/07/2021 | 14:50 | -2.82913 | 57.69946 | -5 | NA | NA | 521 | Near |
| C2.2 | 14/07/2021 | 16:08 | -2.94661 | 57.73386 | -24 | 14.93 | 1.3747 | 4,212 | Middle near |
| C2.3 | 14/07/2021 | 16:25 | -2.94428 | 57.76418 | -34 | 15.02 | 1.1862 | 7,370 | Middle far |
| C2.4 | 15/07/2021 | 14:46 | -2.2303 | 57.69117 | -19 | 14.32 | 9.4317 | 1,115 | Near |
| C2.5 | 19/07/2021 | 13:47 | -2.26489 | 57.72487 | -7 | 15.79 | 1.733 | 721 | Near |
| C2.6 | 19/07/2021 | 14:44 | -3.26542 | 57.78394 | -54 | 14.67 | 0.9296 | 6,571 | Middle near |
| C2.7 | 19/07/2021 | 16:25 | -2.84649 | 57.83421 | -82 | 14.82 | 1.4408 | 14,112 | Far |
| C2.8 | 19/07/2021 | 16:50 | -2.65892 | 57.83317 | -85 | 14.26 | 1.9699 | 15,871 | Far |
| F2.1 | 14/07/2021 | 17:08 | -2.82509 | 57.76437 | -33 | 14.98 | 1.0744 | 6,486 | Middle near |
| F2.2 | 12/07/2021 | 15:20 | -2.54151 | 57.76329 | -104 | 13.49 | 1.0481 | 9,075 | Middle far |
| F2.3 | 15/07/2021 | 16:13 | -2.25834 | 57.76334 | -118 | 14.39 | 1.562 | 7,756 | Middle far |
| S2.1 | 12/07/2021 | 14:44 | -2.5535 | 57.71339 | -50 | 13.25 | 1.0458 | 3,521 | Middle near |
| S2.2 | 14/07/2021 | 10:57 | -2.56572 | 57.72279 | -64 | 14.54 | 0.9909 | 4,383 | Middle near |
| S2.3 | 19/07/2021 | 15:32 | -3.09144 | 57.81815 | -71 | 15.23 | 1.0563 | 14,999 | Far |
| C3.1 | 08/08/2021 | 13:41 | -2.50945 | 57.67346 | -8 | NA | 0.5912 | 488 | Near |
| C3.2 | 10/08/2021 | 17:13 | -2.75408 | 57.76392 | -52 | 15.47 | 0.6513 | 7,301 | Middle far |
| C3.3 | 10/08/2021 | 18:49 | -2.5058 | 57.69665 | -33 | 14.99 | 0.7197 | 2,554 | Middle near |
| C3.4 | 11/08/2021 | 12:16 | -2.4759 | 57.77603 | -97 | 15.27 | 2.0819 | 11,334 | Far |
| C3.5 | 15/08/2021 | 10:04 | -2.81838 | 57.7 | -12 | NA | NA | 468 | Near |
| C3.6 | 15/08/2021 | 13:08 | -2.64304 | 57.68734 | -5 | NA | NA | 236 | Near |
| C3.7 | 15/08/2021 | 16:31 | -2.55517 | 57.67698 | -5 | NA | 1.2377 | 373 | Near |
| F3.1 | 10/08/2021 | 16:57 | -2.82418 | 57.76188 | -32 | 15.7 | 0.5867 | 6,228 | Middle near |
| F3.2 | 10/08/2021 | 18:18 | -2.54113 | 57.76343 | -104 | 15.84 | 0.6061 | 9,095 | Middle far |
| F3.3 | 11/08/2021 | 13:03 | -2.25863 | 57.76345 | -118 | 14.9 | 1.9636 | 7,762 | Middle far |
| S3.1 | 10/08/2021 | 17:56 | -2.57948 | 57.76481 | -88 | 15.86 | 0.6755 | 9,040 | Middle far |
| S3.2 | 15/08/2021 | 10:23 | -2.86333 | 57.70873 | -16 | NA | 1.4816 | 305 | Near |
| C4.1 | 25/09/2021 | 13:51 | -2.63648 | 57.76547 | -82 | 13.39 | 1.5558 | 8,508 | Middle far |
| C4.2 | 28/09/2021 | 11:48 | -2.28578 | 57.69962 | -20 | NA | 4.7475 | 568 | Near |
| C4.3 | 28/09/2021 | 12:34 | -2.16478 | 57.72858 | -56 | 13.62 | 4.5537 | 4,042 | Middle near |
| C4.4 | 28/09/2021 | 13:08 | -2.34827 | 57.72362 | -53 | 13.69 | 4.2259 | 3,958 | Middle near |
| C4.5 | 28/09/2021 | 16:46 | -2.77605 | 57.70878 | -25 | 13.79 | 4.0918 | 1,109 | Near |
| F4.1 | 25/09/2021 | 13:17 | -2.82406 | 57.76314 | -33 | 13.53 | 1.9329 | 6,366 | Middle near |
| F4.2 | 25/09/2021 | 14:22 | -2.54144 | 57.76383 | -104 | 13.17 | 1.4 | 9,135 | Middle far |
| F4.3 | 28/09/2021 | 12:08 | -2.26 | 57.76317 | -118 | 13.34 | 2.0601 | 7,709 | Middle far |
| S4.1 | 04/10/2021 | 14:41 | -2.2043 | 57.7405 | -97 | 13.08 | 2.4202 | 6,579 | Middle near |
| S4.2 | 04/10/2021 | 15:40 | -2.2327 | 57.7788 | -108 | 13.03 | 1.4482 | 9,898 | Middle far |
| S4.3 | 08/10/2021 | 15:46 | -2.1808 | 57.7623 | -49 | 13.03 | 0.9055 | 7,735 | Middle far |

Species of conservation interest were found in the following samples:

Bluefin tuna (*Thunnus thynnus*) – C1.3, C1.4, C1.5, F1.3, S1.2, S1.5, S1.6, C2.1, C2.6, F3.1

European eel (*Anguilla anguilla*) - C2.3, C3.1, C3.2, C3.3, C3.4, C3.5, F3.1, F3.2, S3.1, S3.2. C4.1, C4.3, C4.4

Pink salmon (*Oncorhynchus gorbuscha*) – C1.2, C1.3, C1.5, C1.6, C1.8, F1.2, S1.2, S1.8, C2.1, C2.4, C2.5, C2.6, C2.7, C2.8, F2.1, F2.3, S2.1, S2.2, S2.3, C3.1, C3.2, C3.3, C3.4, C3.5, C3.6, C3.7, S3.2, C4.2, C4.3, C4.4, C4.5

Sowerby’s beaked whale (*Mesoplodon bidens*) – C2.1, C2.2, C2.3, C2.4, C2.5, C2.6, C2.8, F2.1, F2.3, S2.1, S2.2, S2.3, C3.3, F3.1, F3.3, F4.2

Appendix Table A2. List of taxa detected by the different primer sets, MarVer1, MarVer3 and 18S.

| Taxa | Primer set |
| --- | --- |
| Acanthoecaceae | 18S |
| Acartiidae | 18S |
| *Agonus cataphractus* | MarVer3 |
| *Alca torda* | MarVer1, MarVer3 |
| Alciopidae | 18S |
| Ammodytidae | MarVer1, MarVer3 |
| *Anguilla anguilla* | MarVer1, MarVer3 |
| Anisakidae | 18S |
| Archinotodelphyidae | 18S |
| Attheyaceae | 18S |
| Bacillariaceae | 18S |
| *Balaenoptera acutorostrata* | MarVer1, MarVer3 |
| *Balaenoptera physalus* | MarVer1 |
| Batillariidae | 18S |
| *Belone belone* | MarVer3 |
| Beroidae | 18S |
| Bodinidae | 18S |
| Bolinopsidae | 18S |
| Bougainvilliidae | 18S |
| Calanidae | 18S |
| Calciodinellaceae | 18S |
| *Callionymus lyra* | MarVer3 |
| Campanulariidae | 18S |
| Candaciidae | 18S |
| Canthocamptidae | 18S |
| Capitellidae | 18S |
| Catenulaceae | 18S |
| *Centrolabrus exoletus* | MarVer1, MarVer3 |
| Centropagidae | 18S |
| Chaetocerotaceae | 18S |
| Chaetopteridae | 18S |
| *Chirolophis ascanii* | MarVer1 |
| *Ciliata mustela* | MarVer1, MarVer3 |
| *Ciliata septentrionalis* | MarVer1 |
| Cladocorynidae | 18S |
| Clupeidae | MarVer1, MarVer3 |
| Coccolithaceae | 18S |
| Cocconeidaceae | 18S |
| Codonellidae | 18S |
| Codonellopsidae | 18S |
| *Conger conger* | MarVer3 |
| Corethraceae | 18S |
| Corycaeidae | 18S |
| *Crystallogobius linearis* | MarVer1, MarVer3 |
| *Ctenolabrus rupestris* | MarVer1, MarVer3 |
| *Cyclopterus lumpus* | MarVer3 |
| Cylichnidae | 18S |
| Cymatosiraceae | 18S |
| Dasyaceae | 18S |
| *Dicentrarchus* | MarVer1, MarVer3 |
| Dictyochaceae | 18S |
| Dictyotaceae | 18S |
| Didiniidae | 18S |
| Dinophysiaceae | 18S |
| *Diplodus puntazzo* | MarVer1 |
| *Dipturus* | MarVer3 |
| Dotoidae | 18S |
| *Echiichthys vipera* | MarVer3 |
| Ectocarpaceae | 18S |
| *Enchelyopus cimbrius* | MarVer1 |
| Eubranchidae | 18S |
| Eucalanidae | 18S |
| Euchaetidae | 18S |
| Euphausiidae | 18S |
| Euplokamididae | 18S |
| *Eutrigla gurnardus* | MarVer3 |
| Fragilariaceae | 18S |
| *Fratercula arctica* | MarVer3 |
| *Fulmarus glacialis* | MarVer1 |
| Gadidae | 18S |
| *Gaidropsarus mediterraneus* | MarVer3 |
| *Gasterosteus aculeatus* | MarVer1, MarVer3 |
| *Gobiusculus flavescens* | MarVer1 |
| Goniodomataceae | 18S |
| Gonyaulacaceae | 18S |
| *Gymnammodytes semisquamatus* | MarVer3 |
| Gymnodiniaceae | 18S |
| *Halichoerus grypus* | MarVer3 |
| Halocyprididae | 18S |
| Halosphaeraceae | 18S |
| Harpacticidae | 18S |
| Hemiaulaceae | 18S |
| Hemidiscaceae | 18S |
| Hemiselmidaceae | 18S |
| Hepatidae | 18S |
| *Hippoglossoides platessoides* | MarVer1, MarVer3 |
| *Hyperoplus immaculatus* | MarVer1 |
| Isochrysidaceae | 18S |
| *Labrus bergylta* | MarVer1, MarVer3 |
| *Lagenorhynchus albirostris* | MarVer3 |
| *Lamna nasus* | MarVer3 |
| *Lampetra planeri* | MarVer1 |
| *Larus argentatus* | MarVer1, MarVer3 |
| Lauderiaceae | 18S |
| Leptocylindraceae | 18S |
| *Lesueurigobius friesii* | MarVer3 |
| *Leuciscus* | MarVer3 |
| *Leucoraja naevus* | MarVer1, MarVer3 |
| Leucosoleniidae | 18S |
| *Limanda limanda* | MarVer1, MarVer3 |
| Lineidae | 18S |
| *Liparis montagui* | MarVer1, MarVer3 |
| *Lipophrys pholis* | MarVer1, MarVer3 |
| *Lophius piscatorius* | MarVer3 |
| Mamiellaceae | 18S |
| Melosiraceae | 18S |
| Membraniporidae | 18S |
| *Mesoplodon bidens* | MarVer1, MarVer3 |
| Metacylididae | 18S |
| *Micrenophrys lilljeborgii* | MarVer1 |
| *Microchirus* | MarVer1, MarVer3 |
| *Micromesistius* | MarVer3 |
| *Microstomus kitt* | MarVer3 |
| Miliolidae | 18S |
| *Mola mola* | MarVer3 |
| Molgulidae | 18S |
| *Morus serrator* | MarVer3 |
| *Mullus barbatus* | MarVer3 |
| Mycosphaerellaceae | 18S |
| *Myoxocephalus scorpius* | MarVer1, MarVer3 |
| Myrionemataceae | 18S |
| Mytilidae | 18S |
| Naviculaceae | 18S |
| Nephroselmidaceae | 18S |
| Noctilucaceae | 18S |
| *Oncorhynchus* | MarVer1, MarVer3 |
| Opheliidae | 18S |
| Ophiactidae | 18S |
| Pandeidae | 18S |
| Paradiniaceae | 18S |
| Pectinariidae | 18S |
| Pedinellaceae | 18S |
| Peridiniaceae | 18S |
| Phaeocystaceae | 18S |
| *Phalacrocorax carbo* | MarVer1 |
| *Phocoena phocoena* | MarVer1, MarVer3 |
| *Pholis gunnellus* | MarVer1, MarVer3 |
| *Phoxinus phoxinus* | MarVer1, MarVer3 |
| Phyllodocidae | 18S |
| Pleurobrachiidae | 18S |
| Pleuronectidae | 18S |
| Pleurosigmataceae | 18S |
| Podonidae | 18S |
| *Pomatoschistus microps* | MarVer3 |
| *Pomatoschistus minutus* | MarVer1, MarVer3 |
| *Pomatoschistus norvegicus* | MarVer3 |
| *Pomatoschistus pictus* | MarVer3 |
| Prorocentraceae | 18S |
| Prymnesiaceae | 18S |
| Ptychocylididae | 18S |
| Ptychodiscaceae | 18S |
| *Pungitius pungitius* | MarVer3 |
| Pyramimonadaceae | 18S |
| Pyrocystaceae | 18S |
| Pythiaceae | 18S |
| *Raja brachyura* | MarVer1, MarVer3 |
| Ralfsiaceae | 18S |
| *Raniceps raninus* | MarVer3 |
| Rhaphoneidaceae | 18S |
| Rhizophlyctidaceae | 18S |
| Rhizosoleniaceae | 18S |
| Rhodomelaceae | 18S |
| *Rissa tridactyla* | MarVer1, MarVer3 |
| *Salmo* | MarVer1, MarVer3 |
| Saprolegniaceae | 18S |
| *Sardinella longiceps* | MarVer1 |
| Sargassaceae | 18S |
| *Scomber scombrus* | MarVer1, MarVer3 |
| *Scophthalmus rhombus* | MarVer1 |
| *Scyliorhinus canicula* | MarVer1, MarVer3 |
| Scytosiphonaceae | 18S |
| Siphonariidae | 18S |
| Skeletonemataceae | 18S |
| Spathidiidae | 18S |
| *Spinachia spinachia* | MarVer3 |
| Stephanodiscaceae | 18S |
| Striatellaceae | 18S |
| Strombidiidae | 18S |
| *Symphodus melops* | MarVer1, MarVer3 |
| Syndiniaceae | 18S |
| *Syngnathus rostellatus* | MarVer3 |
| Synuraceae | 18S |
| *Taurulus bubalis* | MarVer1, MarVer3 |
| Terebellidae | 18S |
| Tergipedidae | 18S |
| Thalassiosiraceae | 18S |
| *Thorogobius ephippiatus* | MarVer3 |
| Thraustochytriaceae | 18S |
| *Thunnus thynnus* | MarVer1 |
| Tintinnidae | 18S |
| Tomopteridae | 18S |
| *Trachurus* | MarVer3 |
| *Trisopterus esmarkii* | MarVer3 |
| *Trisopterus minutus* | MarVer1, MarVer3 |
| Trochamminidae | 18S |
| *Tursiops truncatus* | MarVer1, MarVer3 |
| Ulmaridae | 18S |
| Ulvaceae | 18S |
| Undellidae | 18S |
| *Uria aalge* | MarVer1, MarVer3 |
| Veneridae | 18S |
| Warnowiaceae | 18S |
| Xystonellidae | 18S |
| *Zeugopterus punctatus* | MarVer3 |
| *Zoarces* | MarVer3 |

Appendix Table A3. Foreign OTUs detected with MarVer1 were compared to native relatives from the same genus or family. Reads were either reassigned to the native relative, removed due to low abundance or identified as a potential invasive.

| **Foreign OTU** | **North Sea relative (shared genus or family)** | **Action** |
| --- | --- | --- |
| *Alectrias benjamini* | *Chirolophis ascanii* | Reassign to native relative |
| *Ammodytes dubius Ammodytes hexapterus Ammodytes personatus* | *Ammodytes marinus Ammodytes tobianus* | All reads combined as *Ammodytes* genus |
| *Anguilla japonica* | *Anguilla anguilla* | Remove as singleton |
| *Chelidonichthys spinosus* | *Eutriglia gurnardus Trigloporus lastoviza* | All reads combined as Triglidae family |
| *Clupea pallasii* | *Clupea harengus* | Reassign to native relative |
| *Diplodus puntazzo* |  | No congeneric in North Sea so potential invasive |
| *Gadus chalcogrammus* | *Gadus morhua Melanogrammus aeglefinus Merlangius merlangus Pollachius pollachius*  *Trisopterus minutus* | All reads combined as Gadidae family |
| *Hippoglossus stenolepsis* | *Hippoglossoides platessoides* | Reassign to native relative |
| *Larus dominicanus* | *Larus argentatus* | Reassign to native relative |
| *Myoxocephalus jaok* | *Myoxocephalus scorpius* | Reassign to native relative |
| *Oncorhynchus clarkii Oncorhynchus mykiss Oncorhynchus nerka* |  | Collapsed to *Oncorhynchus* genus, known invasive |
| *Parahucho perryi Salmo ischchan Salmo labrax* | *Salmo salar Salmo trutta* | Collapsed to *Salmo* genus |
| *Pholis picta* | *Pholis gunnellus* | Reassign to native relative |
| *Sardinella longiceps* | *Clupea harengus Sprattus sprattus* | Distinct from native family members, potential native |
| *Sousa teuszii* | *Tursiops truncatus* | Reassign to native relative |
| *Thunnus maccoyii* | *Thunnus thynnus* | Reassign to native relative |

Appendix Table A4. Foreign OTUs detected with MarVer3 were compared to native relatives from the same genus or family. Reads were either reassigned to the native relative, removed due to low abundance or identified as a potential invasive.

| **Foreign OTU** | **North Sea relative (shared genus or family)** | **Action** |
| --- | --- | --- |
| *Ammodytes americanus Ammodytes hexapterus Ammodytes personatus Hyperoplus lanceolatus* | *Ammodytes marinus Ammodytes Tobianus* | Collapse to ammodytidae family |
| *Anarhichas orientalis* | *Anarhichas lupus* | Remove as <10 reads |
| *Anguilla rostrata* | *Anguilla Anguilla* | Reassign to native relative |
| *Arctogadus glacialis Boreogadus saida* | *Gadus morhua Melanogrammus aeglefinus Merlangius merlangus Pollachius pollachius*  *Pollachius virens Trisopterus esmarkii*  *Trisopterus minutus* | Reassign to native relative, *Gadus morhua* |
| *Clidoderma asperrimum* |  | Remove as singleton |
| *Clupea pallasii* | *Clupea harengus* | Reassign to native relative |
| *Dipturus innominatus*  *Dipturus trachyderma* | *Dipturus batis*  *Dipturus oxyrinchus* | Collapse to *Dipturus* genus |
| *Eleginus gracilis*  *Gadus chalcogrammus*  *Gadus macrocephalus* | *Gadus morhua Melanogrammus aeglefinus Merlangius merlangus Pollachius pollachius*  *Pollachius virens Trisopterus esmarkii*  *Trisopterus minutus* | Remove as <25 reads  Remove as <3 reads  Remove as <3 reads |
| *Glyptocephalus zachirus*  *Hippoglossoides dubius*  *Hippoglossoides elassodon*  *Hippoglossoides stenolepsis*  *Isoptetta isolepsis*  *Lepidopsetta bilineata*  *Lepidopsetta mochigarei*  *Liopsetta pinnifasciata*  *Parophyrs vetulus*  *Platichthys stellatus*  *Prosopium cylindraceum*  *Psettichthys melanostictus*  *Pseudopleuronectes americanus*  *Pseudopleuronectes yokohamae* | *Glytocephalus cynoglossus*  *Hippoglossoides platessoides*  *Limanda limanda*  *Microstomus kitt*  *Pleuronectes flesus*  *Pleuronectes platessa* | Collapse all foreign species and *G. cynoglossus, H. platessoides, P. flesus* and *P. platella* into Pleuronectidae family. Keep *L. limanda* and *M. kitt* separate |
| *Limanda aspera*  *Limanda proboscidea*  *Limanda sakhalinensis* | *Limanda limanda* | Reassign to native relative |
| *Microgadus proximus* | *Gadus morhua* | Reassign to native relative |
| *Myoxocephalus brandtii*  *Myoxocephalus ochotensis*  *Myoxocephalus polyacanthocephalus* | *Myoxocephalus scorpius* | Reassign to native relative |
| *Oncorhynchus clarkii*  *Oncorhynchus gilae*  *Oncorchynchus gorbuscha*  *Oncorhynchus mykiss* |  | Collapse *O. clarkii* and *O. gilae* to *Oncorhynchus* genus but keep *O. gorbuscha* and *O. mykiss* separate. Known invasives. |
| *Phocoena sinus* | *Phocoena phocoena* | Remove as singleton |
| *Pholis laeta* | *Pholis gunnellus* | Reassign to native relative |
| *Pungitius sinensis* | *Pungitius pungitius* | Reassign to native relative |
| *Pusa hispida* | *Halichoerus grypus* | Reassign to native relative |
| *Salmo ischchan*  *Salmo obtusirostris* | *Salmo salar*  *Salmo trutta* | Reassign to native relative, *S. trutta* |
| *Symphodus melamocercus* | *Symphodus melops* | Remove as singleton |
| *Tursiop aduncus* | *Tursiop truncatus* | Reassign to native relative |

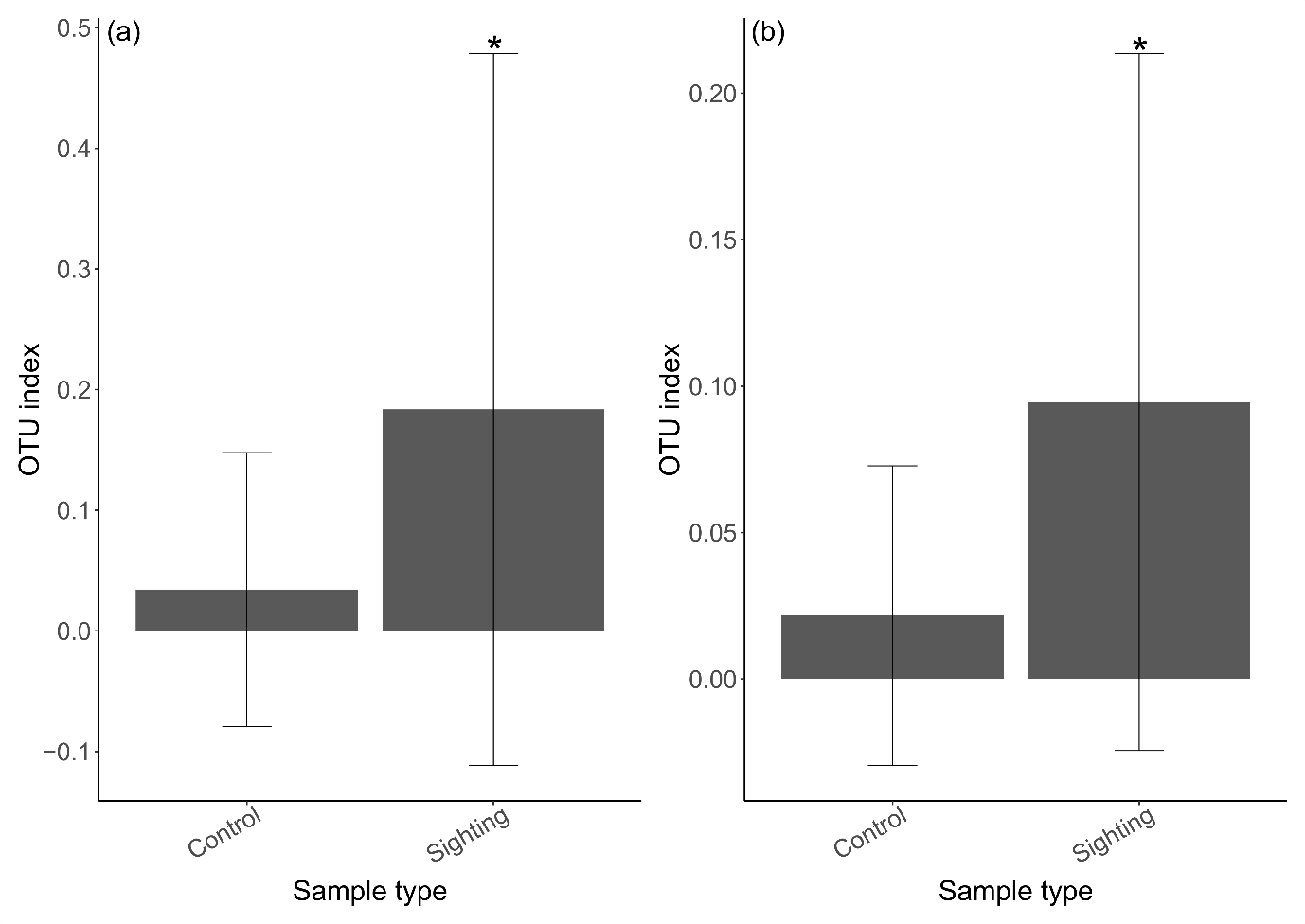

Appendix Figure A1. OTU index for minke whales in control versus sighting samples for (a) MarVer1 and (b) MarVer3. Minke whales were detected in 53 % of sighting samples and 18 % of control samples with MarVer 1 whilst they were detected in 88 % of sighting samples compared to 76 % of control samples with MarVer3. Sighting samples had significantly higher abundances of minke whales compared to control samples for both primer sets (Wilcoxen test, p < 0.05).

Appendix Table A5. Indicator species analysis for vertebrates per trip.

| **Trip** | **OTU** | **A** | **B** | **Stat** | **P value** |
| --- | --- | --- | --- | --- | --- |
| 1 | *Thunnus thynnus* | 0.9946 | 0.3684 | 0.605 | 0.025 |
| 2 | *Mesoploden bidens* | 0.8244 | 0.8571 | 0.841 | 0.015 |
| 2 | *Leuciscus* | 0.9835 | 0.5714 | 0.75 | 0.015 |
| 2 | *Spinachia spinachia* | 0.9751 | 0.5714 | 0.746 | 0.02 |
| 2 | *Diplodus puntazzo* | 1 | 0.4286 | 0.655 | 0.005 |
| 2 | *Fulmarus glacialis* | 1 | 0.2143 | 0.463 | 0.025 |
| 3 | *Oncorhynchus* | 0.9664 | 0.9167 | 0.941 | 0.005 |
| 3 | *Anguilla anguilla* | 0.7431 | 0.75 | 0.747 | 0.005 |
| 3 | *Lesuerigobius friesii* | 0.9983 | 0.4167 | 0.645 | 0.015 |
| 3 | *Mola mola* | 1 | 0.3333 | 0.577 | 0.005 |
| 3 | *Pomatoschistus pictus* | 0.9993 | 0.3333 | 0.577 | 0.02 |
| 4 | *Micrenophrys lilljeborgii* | 0.9947 | 0.6364 | 0.796 | 0.01 |
| 4 | *Mullus barbatus* | 0.9924 | 0.6364 | 0.795 | 0.005 |
| 4 | *Pomatoschistus microps* | 0.9838 | 0.6364 | 0.791 | 0.005 |
| 4 | *Gobiusculus flavescens* | 0.9519 | 0.2727 | 0.51 | 0.02 |
| 4 | *Lophius piscatorius* | 1 | 0.1818 | 0.426 | 0.02 |
| 1+2 | *Balaenoptera acutorostrata* | 0.9328 | 0.8788 | 0.905 | 0.025 |
| 1+2 | *Ciliata septentrionalis* | 0.9167 | 0.7879 | 0.85 | 0.005 |
| 2+3 | *Leucoraja naevus* | 0.974 | 0.7692 | 0.866 | 0.025 |
| 2+3 | *Pleuronectidae* | 0.8742 | 0.8462 | 0.86 | 0.005 |
| 2+3 | *Centrolabrus exoletus* | 0.9824 | 0.7308 | 0.847 | 0.005 |
| 2+3 | *Taurulus bubalis* | 0.8242 | 0.6923 | 0.755 | 0.025 |
| 2+3 | *Dicentrarchus* | 0.994 | 0.5385 | 0.732 | 0.01 |
| 2+3 | *Liparis montagui* | 0.9989 | 0.5 | 0.707 | 0.005 |
| 2+3 | *Zoarces* | 0.9982 | 0.4615 | 0.679 | 0.03 |
| 2+3 | *Chirolophis ascanii* | 0.9996 | 0.3846 | 0.62 | 0.02 |
| 2+4 | *Scyliorhinus canicula* | 0.9587 | 0.68 | 0.807 | 0.015 |
| 3+4 | *Salmo* | 0.9518 | 0.8261 | 0.887 | 0.005 |
| 3+4 | *Trisopterus minutus* | 0.8445 | 0.4783 | 0.636 | 0.01 |
| 1+2+3 | *Gymnammodytes semisquamatus* | 0.996 | 0.9556 | 0.76 | 0.005 |
| 1+2+3 | *Phocoena phocoena* | 0.9999 | 0.8667 | 0.931 | 0.005 |
| 1+2+3 | Ammodytidae | 0.9764 | 0.8222 | 0.896 | 0.005 |
| 1+2+3 | *Uria aalge* | 0.9895 | 0.7778 | 0.877 | 0.03 |
| 1+2+3 | *Ctenolabrus rupestris* | 1 | 0.7556 | 0.869 | 0.005 |
| 1+2+4 | *Ciliata mustela* | 0.9416 | 0.8409 | 0.89 | 0.045 |
| 2+3+4 | *Phoxinus phoxinus* | 0.9998 | 0.6216 | 0.788 | 0.005 |

Appendix Table A6. Indicator species analysis for eukaryotes per trip.

| **Trip** | **OTU** | **A** | **B** | **Stat** | **P value** |
| --- | --- | --- | --- | --- | --- |
| 1 | Tintinnidae | 1.00 | 0.25 | 0.5 | 0.018 |
| 2 | Attheyaceae | 0.9465 | 0.9286 | 0.938 | 0.001 |
| 2 | Rhizophlyctidaceae | 0.9796 | 0.7143 | 0.837 | 0.001 |
| 2 | Metacyclididae | 0.7385 | 0.9286 | 0.828 | 0.002 |
| 2 | Codonellopsidae | 0.9420 | 0.7143 | 0.820 | 0.001 |
| 2 | Campanulariidae | 1 | 0.3571 | 0.598 | 0.002 |
| 2 | Harpacticidae | 1 | 0.3571 | 0.598 | 0.002 |
| 2 | Canthocamptidae | 1 | 0.2143 | 0.463 | 0.036 |
| 3 | Pandeidae | 0.9671 | 0.6667 | 0.803 | 0.001 |
| 3 | Bougainvilliidae | 0.9797 | 0.5833 | 0.756 | 0.001 |
| 3 | Trochamminidae | 0.9263 | 0.5833 | 0.735 | 0.001 |
| 3 | Cylichnidae | 0.9601 | 0.5 | 0.693 | 0.002 |
| 3 | Undellidae | 0.7876 | 0.5833 | 0.678 | 0.002 |
| 3 | Cladocorynidae | 0.9640 | 0.3333 | 0.567 | 0.008 |
| 3 | Dotoidae | 0.9468 | 0.3333 | 0.562 | 0.02 |
| 3 | Molgulidae | 0.8521 | 0.3333 | 0.533 | 0.016 |
| 3 | Corycaeidae | 1 | 0.25 | 0.5 | 0.01 |
| 3 | Dictyotaceae | 1 | 0.25 | 0.5 | 0.013 |
| 4 | Naviculaceae | 0.911 | 1 | 0.954 | 0.001 |
| 4 | Phaeocystaceae | 0.9396 | 0.9091 | 0.924 | 0.001 |
| 4 | Rhaphoneidaceae | 0.9914 | 0.5455 | 0.735 | 0.001 |
| 4 | Veneridae | 0.7542 | 0.3636 | 0.524 | 0.025 |
| 4 | Corethraceae | 0.7495 | 0.3636 | 0.522 | 0.023 |
| 4 | Euchaetidae | 0.9885 | 0.2727 | 0.519 | 0.021 |
| 4 | Eucalanidae | 0.9804 | 0.1818 | 0.422 | 0.047 |
| 1&2 | Didiniidae | 0.9778 | 1 | 0.989 | 0.001 |
| 1&2 | Noctilucaceae | 0.9989 | 0.9706 | 0.985 | 0.001 |
| 1&2 | Spathidiidae | 1 | 0.5882 | 0.767 | 0.001 |
| 1&2 | Xystonellidae | 1 | 0.4706 | 0.686 | 0.002 |
| 1&3 | Bolinopsidae | 0.9274 | 0.9062 | 0.917 | 0.001 |
| 2&3 | Calciodinellaceae | 0.8654 | 1 | 0.930 | 0.001 |
| 2&3 | Pyrocystaceae | 1 | 0.4615 | 0.679 | 0.001 |
| 2&3 | Halosphaeraceae | 0.9906 | 0.4615 | 0.676 | 0.002 |
| 3&4 | Stephanodiscaceae | 0.9308 | 1 | 0.965 | 0.001 |
| 3&4 | Syndiniaceae | 0.9719 | 0.8261 | 0.896 | 0.001 |
| 3&4 | Calanidae | 0.9279 | 0.7826 | 0.852 | 0.001 |
| 3&4 | Lauderiaceae | 0.9941 | 0.4348 | 0.657 | 0.002 |
| 3&4 | Paradiniaceae | 0.89 | 0.3478 | 0.556 | 0.05 |
| 1&2&3 | Prymnesiaceae | 0.9978 | 1 | 0.999 | 0.001 |
| 1&2&3 | Pedinellaceae | 0.9921 | 1 | 0.996 | 0.001 |
| 1&2&3 | Dinophysiaceae | 0.9879 | 1 | 0.994 | 0.001 |
| 1&2&3 | Warnowiaceae | 0.9839 | 1 | 0.992 | 0.001 |
| 1&2&3 | Goniodomataceae | 0.9999 | 0.9565 | 0.978 | 0.001 |
| 1&2&3 | Gonyaulacaceae | 0.999 | 0.913 | 0.955 | 0.001 |
| 1&2&3 | Coccolithaceae | 0.9937 | 0.8913 | 0.941 | 0.001 |
| 1&2&4 | Strombidiidae | 0.9959 | 0.9333 | 0.964 | 0.001 |
| 1&2&4 | Hemiaulaceae | 0.9923 | 0.7556 | 0.866 | 0.002 |
| 1&2&4 | Nephroselmidaceae | 0.9844 | 0.6889 | 0.823 | 0.001 |
| 1&3&4 | Cymatosiraceae | 0.9846 | 0.9535 | 0.969 | 0.002 |
| 1&3&4 | Hemiaulaceae | 0.9475 | 0.9535 | 0.951 | 0.001 |
| 1&3&4 | Euplokamididae | 0.9885 | 0.6279 | 0.788 | 0.004 |
| 1&3&4 | Bodinidae | 0.9653 | 0.5349 | 0.719 | 0.01 |
| 2&3&4 | Fragilariaceae | 0.9778 | 0.5405 | 0.727 | 0.017 |
| 2&3&4 | Ectocaraceae | 0.9738 | 0.5135 | 0.707 | 0.03 |

Appendix Table A7. Indicator species analysis for eukaryotes for distance from shore categories.

| **Trip** | **OTU** | **A** | **B** | **Stat** | **P value** |
| --- | --- | --- | --- | --- | --- |
| Far | Harpacticidae | 0.7433 | 0.3 | 0.472 | 0.035 |
| Near | Ectocarpaceae | 0.80293 | 1 | 0.896 | 0.005 |
| Near | Paradiniaceae | 0.98717 | 0.76923 | 0.871 | 0.005 |
| Near | Ralfsiaceae | 0.95006 | 0.76923 | 0.855 | 0.005 |
| Near | Rhodomelaceae | 1 | 0.69231 | 0.832 | 0.005 |
| Near | Dasyaceae | 0.99224 | 0.69231 | 0.829 | 0.005 |
| Near | Ulvaceae | 0.95142 | 0.61538 | 0.765 | 0.005 |
| Near | Leucosoleniidae | 1 | 0.53846 | 0.734 | 0.005 |
| Near | Molgulidae | 1 | 0.46154 | 0.679 | 0.001 |
| Near | Chaetopteridae | 0.74645 | 0.61538 | 0.678 | 0.01 |
| Near | Siphonariidae | 0.88262 | 0.46154 | 0.638 | 0.015 |
| Near | Catenulaceae | 0.58006 | 0.69231 | 0.634 | 0.015 |
| Near | Ophiactidae | 0.86935 | 0.46154 | 0.633 | 0.005 |
| Near | Lauderiaceae | 0.84958 | 0.46154 | 0.626 | 0.02 |
| Near | Membraniporidae | 1 | 0.38462 | 0.62 | 0.01 |
| Near | Pyrochystaceae | 0.74297 | 0.46154 | 0.586 | 0.01 |
| Near | Archinotodelphyidae | 1 | 0.30769 | 0.555 | 0.015 |
| Near | Scystosiphonaceae | 0.96654 | 0.30769 | 0.545 | 0.015 |
| Near | Myrionemataceae | 0.87352 | 0.30769 | 0.518 | 0.025 |
| Near | Dictyotaceae | 1 | 0.23077 | 0.48 | 0.035 |
| Near | Pectinariidae | 0.98235 | 0.23077 | 0.476 | 0.025 |
| Far + Middle far | Spathidiidae | 0.8088 | 0.5357 | 0.658 | 0.02 |
| Far+middle near+near | Fragilariaceae | 0.977 | 0.5641 | 0.742 | 0.02 |
| Middle far+middle near+near | Stephanodiscaceae | 0.9977 | 0.766 | 0.874 | 0.005 |
| Middle far+middle near+near | Phaeocystaceae | 0.9929 | 0.7021 | 0.835 | 045 |

Appendix Table A8. Indicator species analysis for vertebrates for distance from shore categories.

| **Trip** | **OTU** | **A** | **B** | **Stat** | **P value** |
| --- | --- | --- | --- | --- | --- |
| Far | *Hippoglossoides platessoides* | 0.8783 | 0.5 | 0.663 | 0.005 |
| Near | *Pholis gunnellus* | 0.8521 | 0.9231 | 0.887 | 0.005 |
| Near | *Taurulus bubalis* | 0.9825 | 0.6923 | 0.825 | 0.005 |
| Near | *Labrus bergylta* | 0.9984 | 0.6154 | 0.784 | 0.005 |
| Near | *Zoarces* | 0.9926 | 0.5385 | 0.731 | 0.005 |
| Near | *Liparis montagui* | 0.9913 | 0.5385 | 0.731 | 0.01 |
| Near | *Myoxocephalus scorpius* | 0.9589 | 0.4615 | 0.665 | 0.005 |
| Near | *Mola mola* | 1 | 0.3077 | 0.555 | 0.005 |
| Near | *Pomatoschistus pictus* | 0.9995 | 0.3077 | 0.555 | 0.015 |
| Near | *Gaidropsarus mediterraneus* | 1 | 0.2308 | 0.48 | 0.025 |
| Near | *Thorogobius ephippiatus* | 1 | 0.2308 | 0.48 | 0.015 |
| Far+near | *Limanda limanda* | 0.9326 | 0.8261 | 0.878 | 0.02 |
| Middle near + near | *Tursiops truncatus* | 0.9961 | 0.8214 | 0.905 | 0.005 |
| Middle near +near | *Centrolabrus exoletus* | 0.9843 | 0.5357 | 0.726 | 0.045 |
| Far+middle far+middle near | *Phocoena phocoena* | 0.9759 | 0.8372 | 0.904 | 0.005 |
| Far+middle far+near | Clupeidae | 0.9396 | 0.878 | 0.908 | 0.03 |
| Far+middle near+near | *Gasterosteus aculeatus* | 0.9654 | 0.8421 | 0.902 | 0.03 |
